# Clustered CTCF binding is an evolutionary mechanism to maintain topologically associating domains

**DOI:** 10.1101/668855

**Authors:** Elissavet Kentepozidou, Sarah J Aitken, Christine Feig, Klara Stefflova, Ximena Ibarra-Soria, Duncan T Odom, Maša Roller, Paul Flicek

**Affiliations:** European Molecular Biology Laboratory, European Bioinformatics Institute, Wellcome Genome Campus, Cambridge, CB10 1SD, UK; Cancer Research UK Cambridge Institute, University of Cambridge, Li Ka Shing Centre, Robinson Way, Cambridge, CB2 0RE, UK; Department of Histopathology, Addenbrooke’s Hospital, Cambridge University Hospitals NHS Foundation Trust, Hills Road, Cambridge, CB2 0QQ, UK; German Cancer Research Center (DKFZ), Division Signaling and Functional Genomics, Im Neuenheimer Feld 280, Heidelberg 69120, Germany; Wellcome Sanger Institute, Wellcome Genome Campus, Hinxton, Cambridge, CB10 1SA, UK

## Abstract

CTCF binding contributes to the establishment of higher order genome structure by demarcating the boundaries of large-scale topologically associating domains (TADs). We have carried out an experimental and computational study that exploits the natural genetic variation across five closely related species to assess how CTCF binding patterns stably fixed by evolution in each species contribute to the establishment and evolutionary dynamics of TAD boundaries. We performed CTCF ChIP-seq in multiple mouse species to create genome-wide binding profiles and associated them with TAD boundaries. Our analyses reveal that CTCF binding is maintained at TAD boundaries by an equilibrium of selective constraints and dynamic evolutionary processes. Regardless of their conservation across species, CTCF binding sites at TAD boundaries are subject to stronger sequence and functional constraints compared to other CTCF sites. TAD boundaries frequently harbor rapidly evolving clusters containing both evolutionary old and young CTCF sites as a result of repeated acquisition of new species-specific sites close to conserved ones. The overwhelming majority of clustered CTCF sites colocalize with cohesin and are significantly closer to gene transcription start sites than nonclustered CTCF sites, suggesting that CTCF clusters particularly contribute to cohesin stabilization and transcriptional regulation. Overall, CTCF site clusters are an apparently important feature of CTCF binding evolution that are critical the functional stability of higher order chromatin structure.

## INTRODUCTION

The three-dimensional organization of mammalian genomes comprises distinct structural layers that associate with important functions and range across various scales (Hansen et al., 2018a; Merkenschlager and Nora, 2016; Ruiz-Velasco and Zaugg, 2017). At a scale of tens to hundreds of kilobases, chromatin is partitioned into topologically associating domains (TADs), which are defined as genomic regions with a high frequency of self-interaction, while few or no interactions are observed between neighboring TADs (Dixon et al., 2012; Nora et al., 2012). As a consequence of their insulating structure TADs modulate connections between regulatory elements, such as promoters and enhancers, and thus play an essential role in transcriptional regulation (Mifsud et al., 2015; Nora et al., 2012; Pombo and Dillon, 2015; Schoenfelder et al., 2015; Symmons et al., 2014). TAD structures are reported to be highly conserved across species and cell types (Dixon et al., 2012; Vietri Rudan et al., 2015).

Despite the importance and conservation of TADs, the mechanisms underlying their stability and evolution remain elusive. A large body of evidence supports a model where the CCCTC-binding factor (CTCF), colocalized with the cohesin protein complex, plays a causal role in the formation and maintenance of TADs (Phillips-Cremins et al., 2013; Sofueva et al., 2013; Zuin et al., 2014). CTCF is a ubiquitously expressed zinc-finger protein with a deeply conserved DNA-binding domain (Filippova et al., 1996; Klenova et al., 1993; Moon et al., 2005; Ohlsson et al., 2001). It is responsible for diverse regulatory functions including transcriptional activation and repression as well as promoter and enhancer insulation. Its diverse functions are based on its role in promoting interactions between distant genomic elements by mediating chromatin loop formation (Baniahmad et al., 1990; Lobanenkov et al., 1990; Ong and Corces, 2014). A loop extrusion mechanism of TAD formation has been proposed wherein the cohesin protein complex slides along chromatin forming a growing loop until it meets two CTCF molecules bound with convergent orientation. This architecture then prevents cohesin from sliding further, demarcating the TAD boundaries (Fudenberg et al., 2016; Sanborn et al., 2015). This model explains why these boundaries usually harbor CTCF binding sites. Nevertheless, there are ubiquitous CTCF-bound regions with diverse functions throughout the genome, while only a small fraction of them occur at TAD boundaries (Dixon et al., 2012). This has made it challenging to delineate the precise role of CTCF binding in establishing and stabilizing TAD structures.

Several recent perturbational studies experimentally provide some insights into the role of CTCF in determining local and genome-wide three-dimensional chromatin organization. Local disruption of CTCF binding can lead to abrogation of TAD insulation and formation of ectopic *cis*-regulatory interactions between neighboring TADs (Gómez-Marín et al., 2015; Guo et al., 2015; Nora et al., 2012; Ong and Corces, 2014; Pombo and Dillon, 2015), although TAD structures have been reported to remain intact (Barutcu et al., 2018; Nora et al., 2012; Sanborn et al., 2015). Local TAD disruptions may also lead to disease (Flavahan et al., 2016; Ibn-Salem et al., 2014; Lupiáñez et al., 2015, 2016). Upon acute, transient genome-wide depletion of CTCF there is marked disruption to chromatin loop and TAD structures (Kubo et al., 2017; Nora et al., 2017), but the degree of TAD destabilization remains controversial. The impact of this CTCF-mediated insulation on gene expression remains poorly understood. Indeed, experimental approaches that disrupt CTCF binding remain limited by the fundamental roles of CTCF in development and cell viability.

The binding profiles of CTCF in present-day eukaryotic genomes are shaped by repeated waves of transposable element insertions carrying CTCF binding sequences across mammalian genomes (Bourque et al., 2008; Schmidt et al., 2012; Schwalie et al., 2013; Thybert et al., 2018). Mammalian-conserved sites resulted from ancestral expansions, while recent expansions have established lineage-specific binding patterns. For example, the B2 family of short interspersed nuclear elements (SINEs) active in the mouse-rat ancestor shaped the CTCF binding profile of all Muridae species and specific members of the B2 family remain active in a lineage-specific manner (Bourque et al., 2008; Schmidt et al., 2012; Thybert et al., 2018). The human and macaque genomes also share a large fraction of CTCF-associated transposable elements despite the absence of recent large-scale insertional activity (Schwalie et al., 2013). Moreover, representative mammals share conserved CTCF binding sites at their TAD borders (Dixon et al., 2012; Rao et al., 2014; Vietri Rudan et al., 2015).

The evolutionary history of CTCF binding facilitates a complementary approach to understanding the role of CTCF in TAD stability. Specifically, we can leverage the natural genetic variation between species as opposed to experimental approaches using targeted or systemic CTCF binding disruption. We can thus investigate the consequences of CTCF binding changes stably fixed by evolution as a version of an *in vivo* mutagenesis screen (Heinz et al., 2013). A unique and important advantage of this approach is that the physiological cellular system can be assumed to be in stable and homeostatic equilibrium (Gasch et al., 2016). CTCF is ideally suited to such an evolutionary approach because in each species the CTCF binding profile is composed of substantial numbers of both deeply conserved and evolutionarily recent sites (Schmidt et al., 2012; Thybert et al., 2018).

Here we performed CTCF ChIP-seq in five mouse strains and species, which have similar genomes and transcriptional profiles, to give insight into the establishment and stability of TADs. Our analysis of the genome-wide CTCF binding exploits natural genetic variation between species to assess the evolutionary dynamics of TAD boundary demarcation. We also investigated how local losses of CTCF binding impact gene expression in the neighboring TADs. We revealed that TAD borders are characterized by clusters of both evolutionarily old and young CTCF binding sites. In addition, CTCF bound regions at TAD borders, regardless of age, exhibit increased levels of sequence constraint compared with CTCF binding sites not associated with TAD boundaries. Such clusters are consistent with a model of TAD boundaries in a dynamic equilibrium between selective constraints and active evolutionary processes. As a result, they apparently retain a redundancy of CTCF binding sites that give resilience to the three-dimensional genome structure.

## RESULTS

### *Mus*-conserved CTCF binding sites commonly occur at TAD borders

To investigate the evolution of CTCF binding with respect to the boundaries of topologically associating domains (TADs), we experimentally identified CTCF enriched regions in the livers of five *Mus* species: *Mus musculus domesticus* (C57BL/6), *M. musculus castaneus* (CAST), *M. spretus, M. caroli,* and *M. pahari*. We characterized the conservation level of the identified CTCF binding sites based on whether they are shared by all species (*Mus*-conserved or 5-way), fewer than five species (4-way, 3-way, 2-way) or are species-specific (1-way) (Fig. 1A). The most common categories were the *Mus*-conserved and species-specific CTCF binding sites (Fig. 1A, S1). We found ∼11,000 *Mus*-conserved CTCF binding sites, which made up more than a quarter (∼27%) of the total number of CTCF sites identified in C57BL/6J (Fig. S1). This is consistent with previous observations of high CTCF binding conservation across eutherian mammals, especially compared with other transcription factors such as HNF4A and CEBPA (Kunarso et al., 2010; Schmidt et al., 2010, 2012).

**Figure 1:**
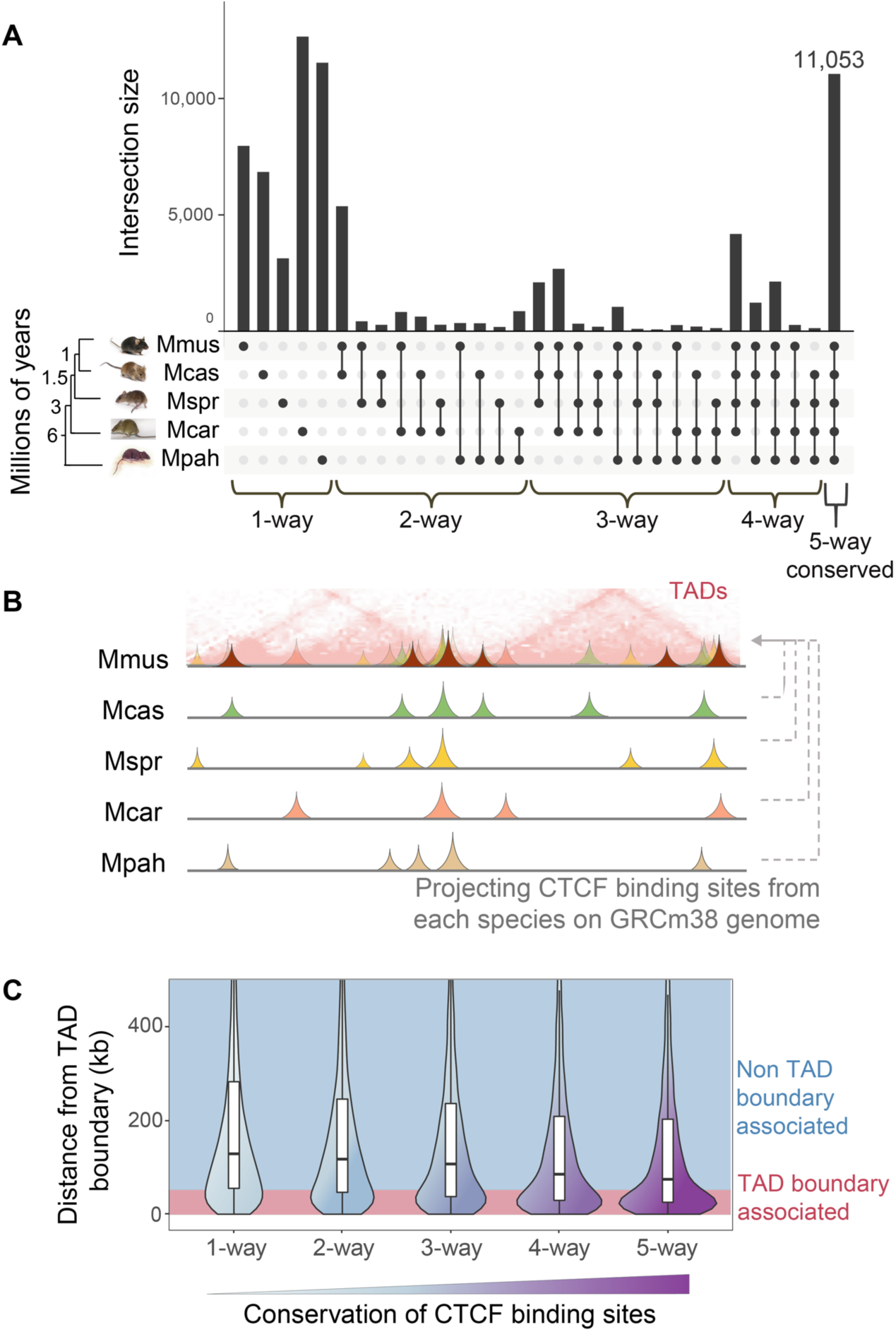
*Mus*-conserved CTCF binding sites commonly occur at TAD borders. (A) Conservation of CTCF binding sites across the five studied *Mus* species. Conservation levels, i.e. the number of species CTCF sites are shared in, are noted at the bottom of the panel (phylogenetic distances are from Thybert et al., 2018). (B) Graphical representation of using orthologous alignments of the CTCF sites identified in each *Mus* species to project them on the genome of C57BL/6J (Mmus, GRCm38) where TADs are available. (C) Distances of CTCF sites with different conservation levels to their closest TAD boundary. CTCF sites with a distance ≤50kb are considered TAD-boundary associated, while sites with a distance >50kb are referred to as non-TAD-boundary associated.

We then intersected the CTCF binding profiles with TAD borders identified using Hi-C in C57BL/6J liver (Vietri Rudan et al., 2015). We projected the CTCF sites identified in each of the five *Mus* species onto the C57BL/6J genome assembly (GRCm38/mm10) (Fig. 1B). After grouping all the CTCF sites by conservation level, we measured the distance from each CTCF site to its closest TAD boundary. Based on this distance and the resolution of the TAD map used, we distinguished between TAD-boundary-associated (*d* ≤ 50kb) and non-TAD-boundary-associated CTCF binding sites (*d* > 50kb). We observed that, although CTCF sites of all conservation levels associate with TAD boundaries, more highly conserved CTCF sites were, on average, located closer to TAD boundaries (Fig. 1C). Overall, 41% of the *Mus*-conserved CTCF sites, as compared to 23% of species-specific sites, were found to lie within 50kb of TAD boundaries (Fig. S2). Our finding of a progressive evolutionary trend between TAD boundaries and CTCF binding conservation, even among closely related species, supports previous reports that shared human-mouse (Rao et al., 2014) and mouse-dog binding sites overlap with the boundaries of TADs (Vietri Rudan et al., 2015).

Shifting the perspective from CTCF bound regions to TAD boundaries, we found that the majority of TAD borders overlap with highly-conserved CTCF binding sites. Nevertheless, a small fraction of the boundaries did not harbor any *Mus*-conserved CTCF binding events. In particular, twelve percent had CTCF sites conserved only in one, two or three out of the five studied *Mus* species (Fig. S3). Furthermore, nearly 5% of TAD boundaries apparently do not overlap with any CTCF occupancy (Fig. S3). One potential interpretation is that, although the connection between CTCF binding and TAD boundaries was consistently observed, it may not strictly necessary feature for demarcation of TAD boundaries as suggested by Hansen et al., 2018a.

In summary, the majority of CTCF binding sites are conserved across five mouse species. Moreover, 41% of *Mus*-conserved CTCF binding sites were associated with a TAD boundary, while the vast majority (>95%) of all TAD boundaries have at least one CTCF binding site.

### CTCF binding sites at TAD boundaries are under strong evolutionary constraint

To investigate the role of TAD boundary association in shaping the characteristics of CTCF binding sites we first assessed the relationship among CTCF conservation level, TAD boundary association, and CTCF motif strength. Specifically, we identified CTCF motifs from our ChIP-seq peaks and calculated their binding affinity (see Methods). CTCF is known to bind to a 33/34 base pair region of the genome consisting of a primary sequence motif (M1) and a shorter secondary motif (M2) (Schmidt et al., 2012). We found that overall binding affinity was significantly greater for boundary-associated CTCF sites compared to non-boundary-associated sites (Mann-Whitney U test, *p* < 2.2e-16) (Fig. 2A). We asked whether this increase in affinity is driven by the fact that many *Mus-*conserved CTCF sites overlap with TAD boundaries. Although motif binding affinity increased with the CTCF binding site conservation level, TAD-boundary-associated CTCF binding sites consistently had greater binding affinity than non-boundary-associated sites (Mann-Whitney U tests between TAD-boundary-associated and non-TAD-boundary-associated sites: *p*_5-way_= 3.9e-11, *p*_4-way_= 5.2e-13, *p*_3-way_= 6.1e-07, *p*_2-way_= 0.06, *p*_1-way_= 0.001) (Fig. 2B). In addition, we confirmed that, independent of conservation level, CTCF binding sites at TAD borders show higher ChIP enrichment than non-TAD-boundary-associated CTCF sites, (Fig. 2C, 2D) consistent with the stronger predicted affinity for CTCF. Overall, our results give new insight into the observation that mammalian-conserved CTCF sites have higher motif affinity than species-specific sites (Schmidt et al., 2012; Vietri Rudan et al., 2015). Importantly, for all CTCF binding sites, including species-specific ones, proximity to a TAD boundary was associated with an increase in binding affinity (Fig. 2B, 2D). This implies that CTCF binding motifs at TAD boundaries may be under stronger selective constraint than the motif sequences of non-TAD-boundary-associated CTCF peaks.

**Figure 2:**
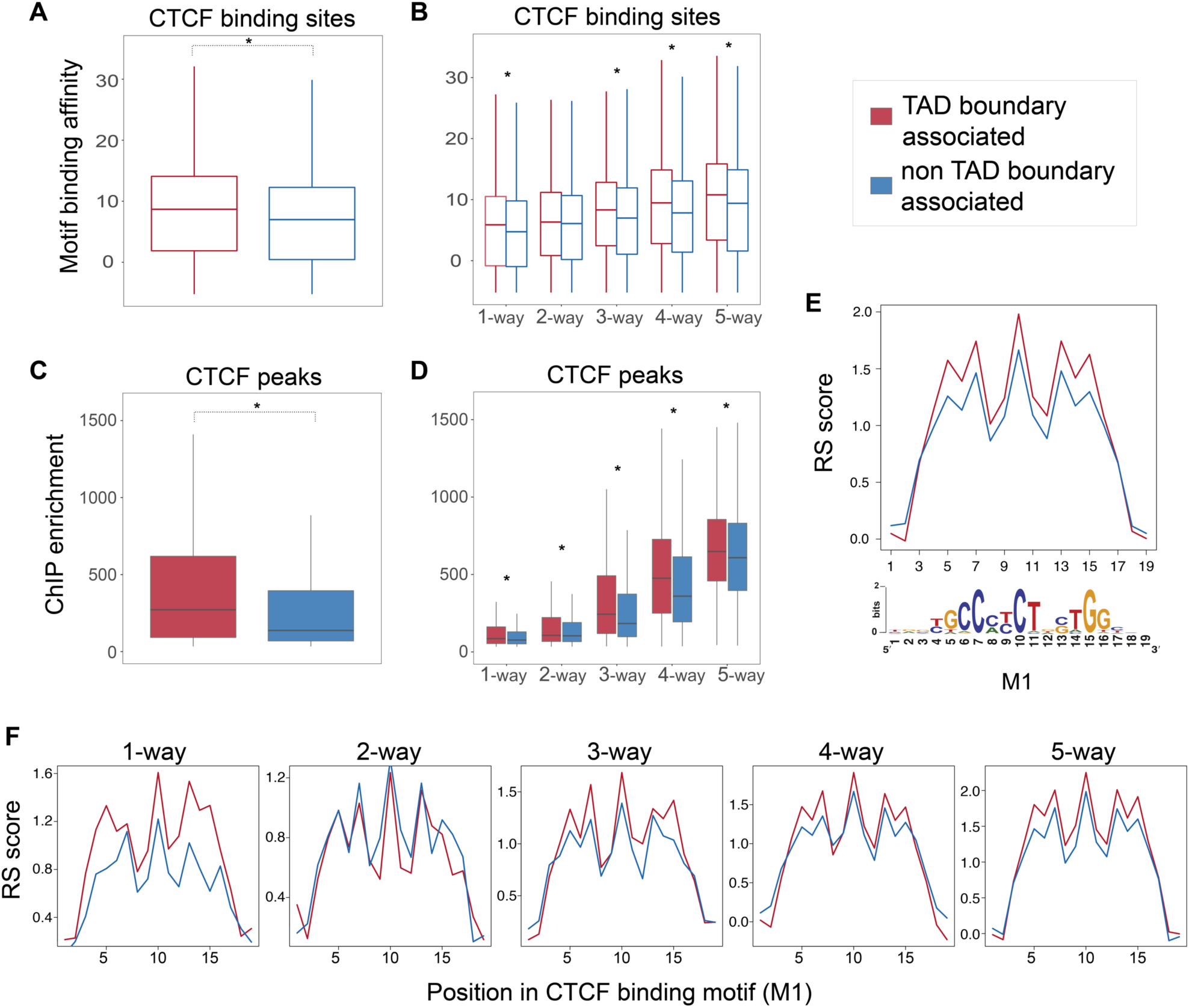
CTCF binding sites at TAD boundaries are subjected to stronger evolutionary constraints, (A) CTCF-bound sites at TAD boundaries contain motifs with higher binding affinity for CTCF than non-TAD-boundary-associated sites (Mann-Whitney U test: p-value < 2.2e-10). (B) Although the binding affinity of CTCF sites is generally proportional to the conservation level of the site (how many species it is shared by), CTCF sites at TAD boundaries have stronger binding affinity than non-TAD-boundary-associated sites, independent of their conservation level (Mann-Whitney U tests between TAD-boundary-associated and non-TAD-boundary-associated sites: *p*_1-way_= 0.001, *p*_2-way_= 0.06, *p*_3-way_= 6.1e-07, *p*_4-way_= 5.2e-13, *p*_5-way_= 3.9e-11). (C) TAD-boundary-associated CTCF peaks display higher ChIP enrichment scores, as calculated by MACS, than non-TAD-boundary-associated peaks (Mann-Whitney U test: p-value < 2.2e-10). (D) TAD-boundary-associated CTCF peaks, at every conservation level, display stronger ChIP enrichment than non-TAD-boundary-associated peaks (Mann-Whitney U tests: *p*_1-way_ < 2.2e-16, *p*_2-way_= 0.002316, *p*_3-way_< 2.2e-16, *p*_4-way_< 2.2e-16, *p*_5-way_= 2.047e-12). (E) The most information-rich bases of the primary CTCF M1 motif at TAD boundaries display higher rejected substitution (RS) scores compared to non-TAD-boundary-associated motifs. The bottom panel shows the position weight matrix of the CTCF M1 motif from Schmidt et al., 2012. (F) The observation in *(E)* is independent of the conservation level of the CTCF sites, as shown for subsets of sites at each conservation level.

To investigate this hypothesis, we explored evolutionary sequence constraint of the CTCF binding motif itself. We estimated sequence constraint by measuring the rejected substitution rate (RS score) at each position of every 19 base-long primary CTCF binding motif (M1) and compared the score between (a) TAD-boundary-associated and (b) non-TAD-boundary-associated regions (Fig. 2E, 2F). RS score is a measure of sequence constraint and reflects the number of base substitutions that were rejected at a specific genomic position as a result of purifying selection, compared to the number of substitutions that would have occurred if the sequence was evolving under neutral selection (Cooper, 2005). We found that the M1 motif in TAD-boundary-associated sites displayed higher RS scores compared to the motifs of non-TAD-boundary-associated sites (Fig. 2E). We further compared the mean RS score per base between the two categories for CTCF sites at every conservation level and confirmed the generality of this observation (Fig. 2F). We also established that this observation was not caused by an enrichment of specific motif instances at TAD boundaries (Fig. S4).

Taken together, CTCF binding sites at TAD boundaries are subject to stronger evolutionary constraints than the CTCF binding sites that are located further away, and that this relationship is independent of evolutionary origin of the site.

### LINEs and LINE-derived CTCF sites are under-represented at TAD boundaries

Having observed that localization of CTCF sites at TAD boundaries affects their sequence and functional conservation, we questioned whether CTCF binding near TAD boundaries appears to evolve by specific mechanisms. Previous results demonstrate that the binding profile of CTCF in eukaryotic genomes is, to a large extent, the consequence of repeat element expansion (Bourque et al., 2008; Schmidt et al., 2012; Sundaram et al., 2014; Thybert et al., 2018). We searched for potential differences in the transposon classes that drive CTCF binding expansion at TAD boundaries compared to the whole genome. We grouped the CTCF sites based on whether they locate at TAD boundaries or not, and for each group we calculated the number of CTCF peak centers that were embedded in SINEs, long terminal repeats (LTRs), long interspersed nuclear elements (LINEs), and DNA transposons. As expected, the greatest fraction of CTCF sites in both categories were found to be SINE-derived (Fig. 3A) (Bourque et al., 2008). The fraction of SINE-derived CTCF sites at TAD borders was slightly, but not significantly, larger than in the rest of the genome (χ^2^ test without Yates correction: *p* = 0.01), implying that SINEs may have uniform potential to establish a CTCF site at both TAD boundaries and other genomic regions. Similarly, CTCF sites of LTR origin did not show significant differences between the two categories (χ^2^: *p* = 0.015). In contrast, the relative proportion of DNA transposon-derived CTCF sites was increased at TAD boundaries (χ^2^: *p* = 0.0003) but accounted for less than 3% of the TEs that contribute to CTCF binding (Fig. 3A). The depletion of LINE-derived CTCF binding sites at TAD boundaries compared to the background genome was the most striking difference (χ^2^: *p* = 3.147e-15; Fig. 3A) suggesting that CTCF binding site formation via LINE expansion is significantly less common at TAD borders than genome-wide.

**Figure 3.**
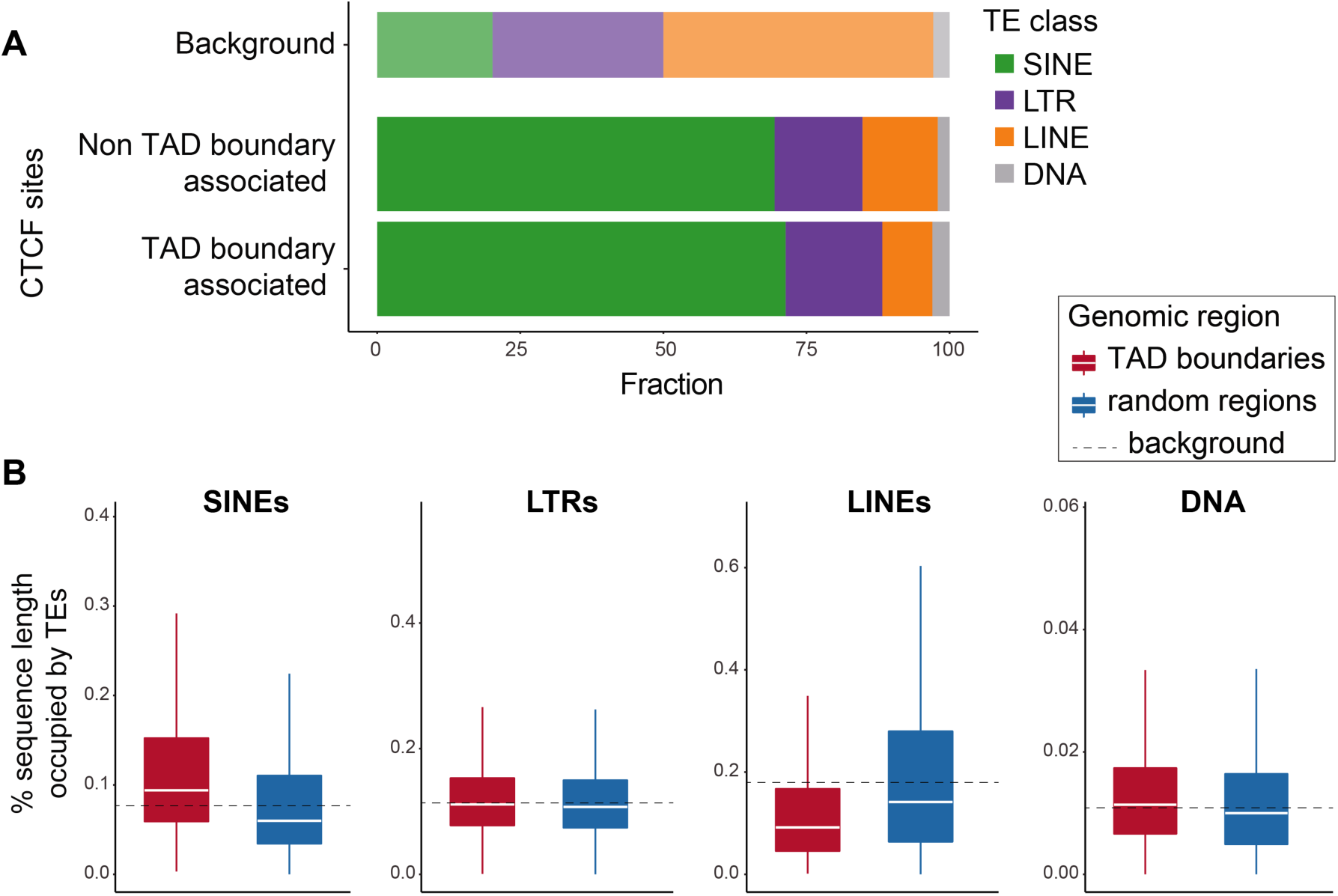
Representation of TE classes and their association with CTCF binding sites differs between TAD boundaries and other genomic regions. (A) Fractions of TAD-boundary-associated versus non-TAD-boundary-associated CTCF binding sites that are embedded in different TE classes. LINE-embedded CTCF-sites are under-represented at TAD boundaries (χ^2^ test without Yates correction: *p* = 3.12e-15), while DNA-transposon-embedded CTCF sites are over-represented (χ^2^ test: *p* = 0.0003), although accounting for just 3% of the TAD-boundary-associated sites. SINE-derived CTCF sites (χ^2^ test: *p* = 0.01) and LTR-associated CTCF sites (χ^2^ test: *p* = 0.015) show no significant differences between the two categories. The top bar shows the percentage of the C57BL/6J genome sequence that corresponds to each TE class, for reference. (B) Fraction of sequence length of TAD boundary regions (TAD boundary +/-50kb) occupied by each TE class, compared to random genomic regions of equal length. SINE sequences are significantly over-represented (Mann-Whitney U test: *p* < 2.2e-16), while LINEs are significantly depleted at TAD boundaries (*p* < 2.2e-16). DNA transposons are slightly, but significantly, enriched at TAD borders (*p=* 9.72e-14), although they account for only 1% of the sequences of the studied regions on average. Representation of LTR sequences shows no significant difference between TAD boundaries and random genomic regions (*p=* 0.005; significance threshold: 0.001).

We further assessed the representation of SINE, LTR, LINE, and DNA transposon sequences around TAD boundaries, independent of whether they carry CTCF binding sites. In particular, we determined the fraction of the 100kb TAD border regions occupied by different transposon classes and compared these with random genomic regions of similar size and distribution. SINE sequences were significantly enriched at TAD boundaries (Mann-Whitney U test: *p* < 2.2e-16; Fig. 3B) (Dixon et al., 2012). The fraction of LTR-derived sequences at TAD boundaries was only marginally higher than random genomic regions (*p*=0.005), and the fraction of DNA transposon sequences was also slightly higher at TAD borders (*p* = 9.72e-14; Fig. 3B). In contrast, LINE sequences were significantly under-represented at TAD boundaries, compared to random genomic regions (Mann-Whitney U test: *p* < 2.2e-16; Fig. 3B), suggesting that TAD boundaries are depleted of LINEs, which may explain why LINE-derived CTCF sites appear under-represented at TAD boundaries (Fig. 3A). Considering the characteristic length of LINE elements, this observation potentially indicates that the insertion of long sequences such as LINEs is negatively selected at TAD borders. This result is complementary to recent reports of selection against long sequence deletions at the functional regions of TAD boundaries (Fudenberg and Pollard, 2019). Moreover, it extends our previous observations and reinforces the hypothesis that in addition to TAD-boundary-associated CTCF sites being subjected to stronger sequence and functional constrains, TAD boundary regions as a whole are under stronger evolutionary pressure (Fudenberg and Pollard, 2019).

### TAD borders harbor clusters of conserved and nonconserved CTCF binding sites

To gain further insight into the architecture of TAD boundaries we investigated the organization of CTCF binding sites within them. In particular, we examined how the density of CTCF binding sites is related to distance from the TAD boundary. By grouping the CTCF binding sites based on conservation level we observed that, as expected, TAD borders were highly enriched for conserved CTCF binding events (Fig. 4A). However, species-specific CTCF binding sites were, surprisingly, also enriched at TAD boundaries (Fig. 4A). Thus, TAD boundaries harbor both numerous conserved CTCF binding sites and a high concentration of species-specific CTCF sites. Additionally, TAD-boundary-associated sites were consistently close to a neighboring site (*median distance* ≈ 5.3kb-5.9kb) regardless of their conservation level (Fig. 4B). In contrast, CTCF binding sites not associated with a TAD boundary were further apart from each other (Mann-Whitney U test: *p* < 2.2e-16) and the median distance to their closest neighboring site was dependent on conservation level: 7kb for 5-way conserved sites to 10.5kb for species-specific sites (Fig. 4B).

**Figure 4:**
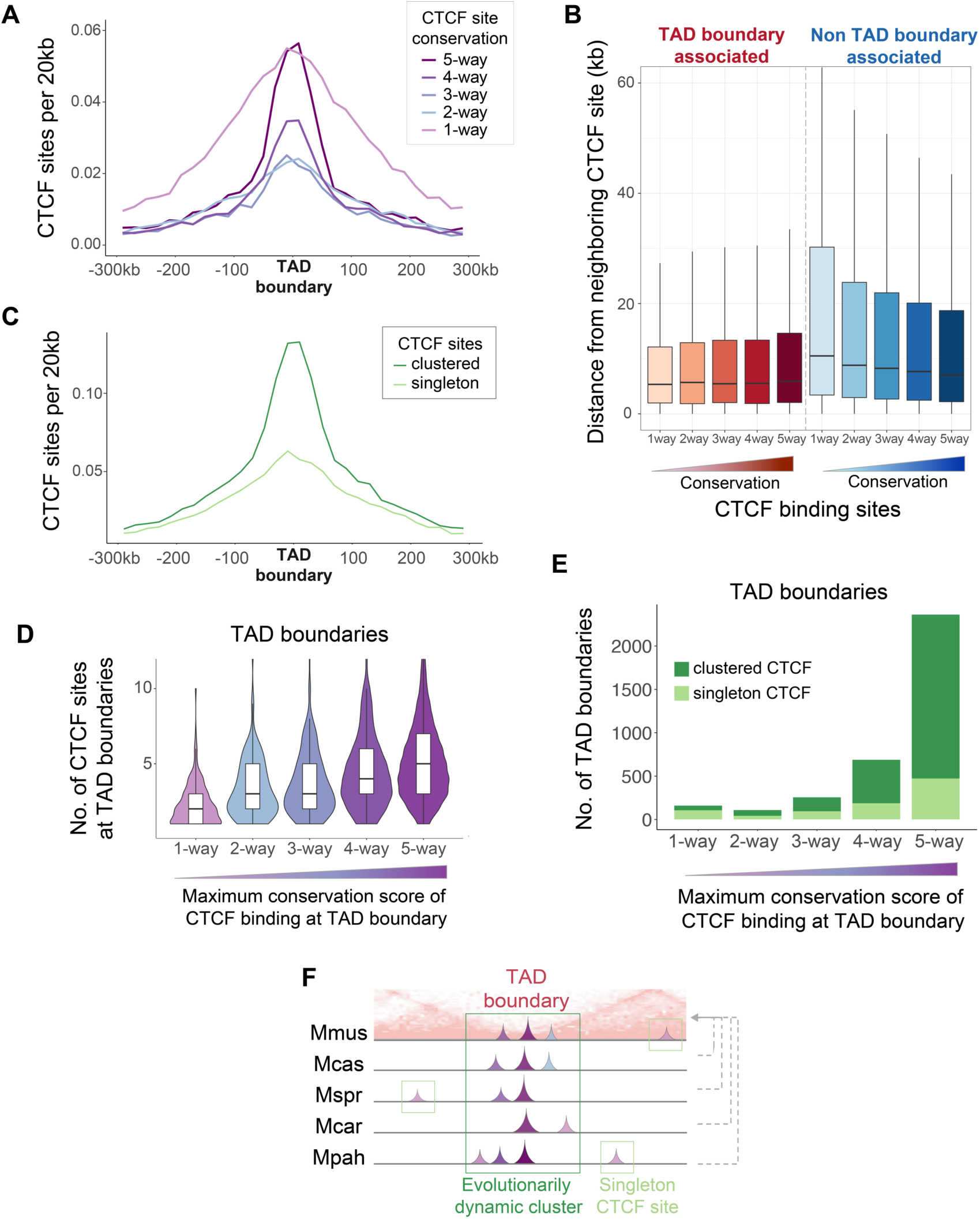
TAD boundaries harbor clusters of both conserved and divergent CTCF binding sites. (A) Both *Mus*-conserved and species-specific CTCF binding sites are highly enriched around TAD boundaries. CTCF sites shared by 2-4 species are also enriched around TAD boundaries. (B) TAD-boundary-associated sites lie significantly closer to each other compared to non-TAD-boundary-associated CTCF sites (Mann-Whitney U test: *p* < 2.2e-16). (C) CTCF binding sites that belong to a cluster (“clustered”) are more enriched at TAD boundaries than singleton CTCF sites. (D) The violin plots correspond to TAD boundaries categorized according to the maximum conservation level of CTCF binding they contain. Each violin plot shows the distribution of the total number of CTCF sites that occur at the TAD boundaries in the category. TAD boundaries with at least one *Mus*-conserved site (right-most violin plot) also have a higher number of CTCF sites overall (higher redundancy). In contrast, TAD boundaries that do not contain any species-conserved CTCF sites (left-most violin plot) have much lower numbers of CTCF binding sites. There is a progressive association between presence of individual conserved CTCF sites with higher abundance of CTCF sites. (E) The bars correspond to TAD boundaries categorized according to the maximum conservation level of CTCF binding they contain. Dark green demarcates TAD boundaries with clustered CTCF sites; light green shows TAD boundaries with only singleton sites. TAD boundaries that harbor species-conserved CTCF sites also contain CTCF site clusters. (F) Schematic representation of evolutionarily dynamic clusters of CTCF sites that commonly occur at TAD boundaries. TAD borders usually have at least one 5-way conserved CTCF site that is clustered with other sites of lower conservation, including species-specific ones. These CTCF clusters preserve CTCF binding potential at TAD boundaries.

We asked whether TAD borders have a specific structure of CTCF sites by investigating potential ancestral clusters from the full set of CTCF binding sites projected to the C57BL/6J genome (*n* = 56,625; Fig. 1B). We defined a CTCF cluster as a group of at least two CTCF binding sites that are each less than 10kb apart on the genome. After clustering we found that 23,232 (43%) sites were singletons whereas 32,393 (57%) were part of 11,507 clusters. Interestingly, we observed that the CTCF sites belonging to a cluster were significantly enriched at TAD borders than singleton CTCF sites (Fig. 4C). This finding strongly implies that clusters of CTCF binding sites are a fundamental architectural structure of TAD boundaries.

To further characterize the CTCF binding clusters at TAD borders, we asked how features such as redundancy, clustering, and presence of both conserved and nonconserved binding events lying in close proximity are associated with each other. We found that TAD borders with at least one 5-way conserved CTCF site contained both a higher number of CTCF sites overall (Fig. 4D) that mainly belong to clusters (Fig. 4E). This shows that *Mus-*conserved CTCF sites at TAD boundaries usually form clusters with other, more recently evolved CTCF sites (Fig. 4F).

We questioned whether this phenomenon is solely a characteristic of TAD boundaries or is it also found in other part parts of the genome. We identified 5-way conserved CTCF sites that were not associated with TAD boundaries (selected as *d* > 80kb from the TAD border to ensure the entire cluster would be *d* > 50kb) and inspected the CTCF binding profile around them. We observed that additional CTCF sites of various conservation levels, including high numbers of species-specific CTCF sites, were generally accumulated around these *Mus*-conserved sites (Fig. S5). Overall, *Mus-*conserved CTCF binding events are usually part of CTCF binding clusters, rather than appearing as singleton sites. Moreover, although the clusters are apparently stably anchored at 5-way CTCF sites, the cluster as a whole seems to be evolving dynamically, allowing for integration of many evolutionarily younger lineage-specific sites.

Finally, we investigated whether the evolutionary characteristics of clustered CTCF binding across the five species were recapitulated when looking at a single species. We confirmed the enrichment of C57BL/6J CTCF sites of any conservation level at TAD boundaries (Fig. S6A) and that clustered CTCF sites in C57BL/6J were also more highly enriched at TAD boundaries than singleton CTCF sites (Fig. S6B), as observed in all *Mus* species (Fig. 4A, 4C). Moreover, we found that half of C57BL/6J CTCF binding sites were clustered, similar to the full set of *Mus* CTCF binding regions (Fig. S6C). We also found that the conservation of whole clusters of CTCF sites in C57BL/6J was similar to that of individual CTCF binding sites (Fig. S6D). This implies that clusters of CTCF sites are evolving under selective pressure similar to that underlying the conservation of individual CTCF binding sites.

In summary, clusters of CTCF binding sites of all conservation levels are a common characteristic of TAD boundaries maintained by dynamic evolutionary processes with species-specific sites playing a prominent role. In addition, CTCF clusters with similar characteristics can also be found distant to TAD borders suggesting a broader role in genome function.

### Clusters of CTCF binding sites colocalize with cohesin and regulate gene expression

To gain further insight into possible additional functional roles of CTCF binding site clusters, we performed ChIP-seq for the cohesin subunit RAD21 in C57BL/6J. CTCF is known to interact with cohesin to form chromatin loops (Ong and Corces, 2014; Parelho et al., 2008; Rubio et al., 2008; Stedman et al., 2008; Wendt et al., 2008; Xiao et al., 2011). To control for the longer genomic regions spanned by CTCF clusters we extended the genomic intervals around the singleton CTCF sites such that the mean of their length distribution was equal to that of the CTCF site clusters (Fig. S7). We found that CTCF site clusters were significantly more likely to overlap with regions enriched for RAD21; 93% compared with only 69% for singleton CTCF sites (χ^2^ test, *p* <2.2e-16) (Fig. 5A). This suggests that clusters of closely located CTCF binding sites help stabilize cohesin and may represent anchors of chromatin loops or TAD boundaries.

**Figure 5:**
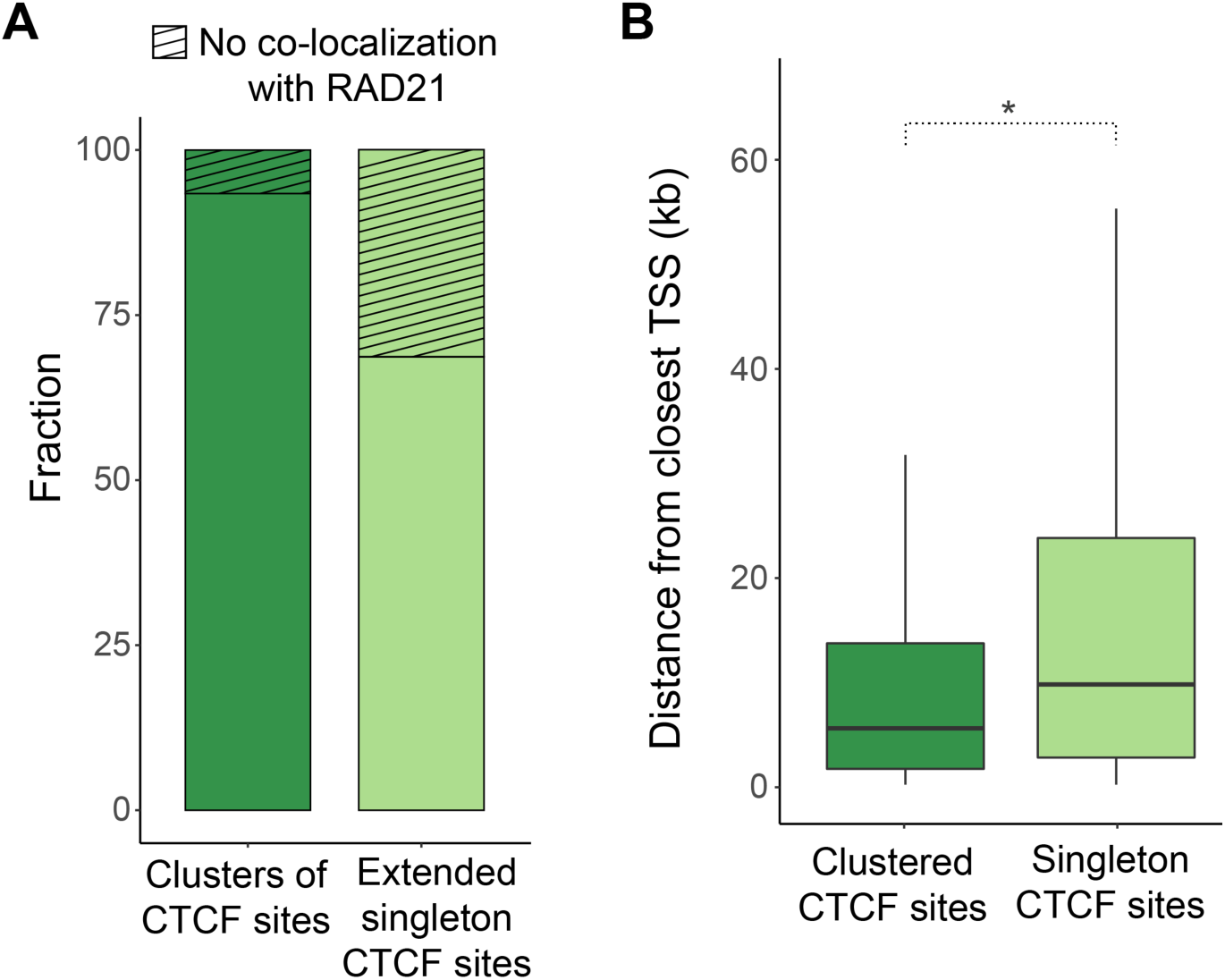
Clustered CTCF sites overlap more frequently with cohesin and locate closer to genes, compared to singleton CTCF binding sites. (A) 93.7% of the clusters of CTCF binding sites colocalize with the cohesin subunit RAD21, while the respective fraction of extended singleton CTCF sites is 69% (χ^2^ test: *p* < 2.2e-16). The singleton CTCF binding regions were extended by a few kilobases prior to intersection with RAD21 binding regions to ensure the mean of their length distribution is equal to the mean length distribution of clusters of CTCF sites. (B) CTCF sites that belong to clusters (clustered) are located closer to gene TSSs (Median distance = 5.3kb) than singleton CTCF sites (Median distance = 10.9kb) (Mann-Whitney U test: *p* < 2.2e-16).

CTCF is also known to bind near gene promoters (Chen et al., 2012). We measured the distance of each CTCF site belonging to a cluster to the nearest transcription start site (TSS) and compared this distribution to the corresponding distances for singleton CTCF sites. We found that CTCF sites belonging to a cluster are generally located significantly closer to TSSs (Median distance = 5.3kb) than singleton CTCF sites (Median distance = 10.9kb) (Mann-Whitney U test, *p* < 2.2e −16; Fig. 5B) which suggests that clusters of CTCF sites may also play an integral role in regulating gene expression.

### The insulating function of CTCF at TAD boundaries is robust to species-specific loss of conserved binding events

CTCF binding sites at TAD boundaries are thought to enhance contact insulation between regulatory elements of adjacent TADs (Schoenfelder et al., 2015) and therefore their disruption can lead to local ectopic interactions between promoters and enhancers (Flavahan et al., 2016; Guo et al., 2015; Nora et al., 2012). However, the impact of such disruptions on local gene expression has not been systematically investigated. Here, we took advantage of natural genetic variation in closely related mouse species and our own CTCF binding data to study the effect of CTCF binding site loss in a model fixed by evolution. This approach offers significant advantages over many other experimental approaches, such as disruption of specific CTCF sites (Barutcu et al., 2018; Guo et al., 2015; Lupiáñez et al., 2015; Nora et al., 2012), haploinsufficiency models (Kemp et al., 2014), or transient acute depletion systems (Kubo et al., 2017; Nora et al., 2017) in which there is global disruption of cellular equilibrium.

We investigated the instances at TAD boundaries where a CTCF binding event was conserved in all but one of the five study species. We estimated the impact of these changes on the expression of proximal genes using RNA sequencing (RNA-seq) in C57BL/6J, CAST, and *M. caroli*. First, we identified either CAST-specific (Fig. 6A) or *M. caroli-*specific losses of individual CTCF binding events at TAD boundaries (Fig. 6D). For each of these lost CTCF sites, we found the closest upstream and the closest downstream one-to-one orthologous gene in all three species (Fig. 6A, 6D) and calculated the relative gene expression of this gene pair (expressed as *log*_*2*_ *fold-change*) in each of the species (see Methods). We then compared these relative expression patterns among the three species.

**Figure 6:**
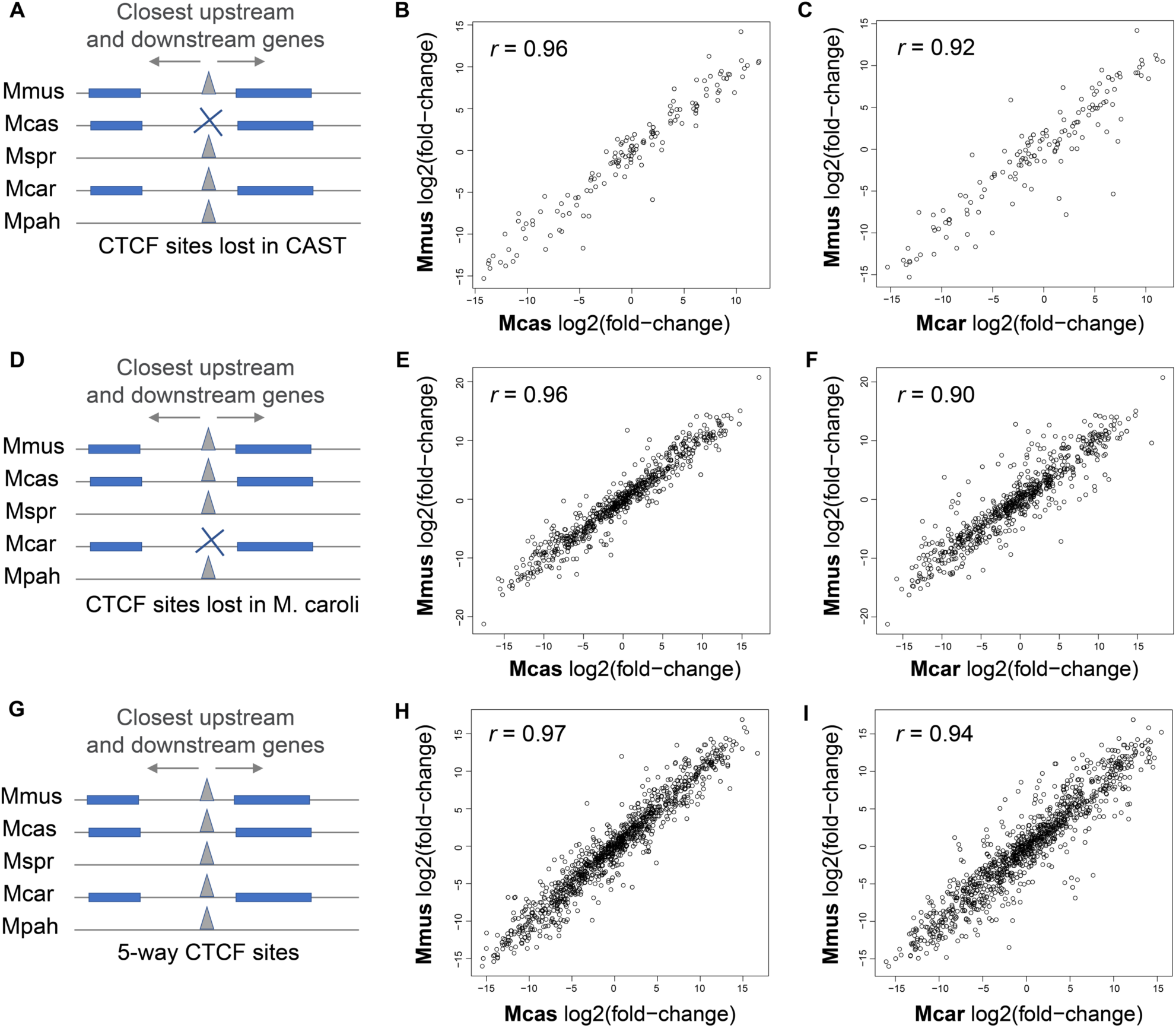
Gene expression patterns around TAD boundaries are robust to local species-specific losses of individual CTCF sites. (A) We identified *M. musculus castaneus* (CAST)*-*specific CTCF site losses at TAD boundaries and estimated the gene expression patterns around them, by calculating the log_2_(fold-change) between the closest downstream to the closest upstream gene. (B, C) Comparisons of log_2_(fold change) values of gene pairs flanking the CAST-specific losses of CTCF sites between C57BL/6J and CAST, with inconsistent CTCF binding, as well as between C57BL/6J and *M. caroli,* with consistent CTCF binding. Only genes that have a one-to-one orthologous relationship and similar gene lengths among C57BL/6J, CAST, and *M. caroli* were used. (D) *M. caroli-*specific CTCF site losses at TAD boundaries and estimated the gene expression patterns around them, with calculated log_2_(fold change) between the closest downstream to the closest upstream gene. (E, F) Comparisons of log_2_(fold change) values of gene pairs flanking the *M. caroli*-specific losses of CTCF sites between C57BL/6J and CAST, with consistent CTCF binding, as well as between C57BL/6J and *M. caroli,* with inconsistent CTCF binding. (G) For reference, *Mus-*conserved CTCF sites and calculated gene expression patterns around them with computed the log_2_(fold change) of the closest downstream to the closest upstream gene in each of the species. (H, I) Comparisons of log2(fold-change) values of gene pairs flanking the examined *Mus-*conserved CTCF sites between C57BL/6J and CAST, as well as between C57BL/6J and *M. caroli.*

We found no impact on insulating function due to species-specific losses of individual CTCF binding events at TAD borders (Fig. 6B, 6C, 6E, 6F, 6H, 6I). This suggests that expression patterns of genes at the borders of TADs are robust to the losses of individual CTCF binding even in cases where the binding event is preserved in multiple other closely related species. We propose that the observed CTCF clusters, which may function interchangeably or additively, contribute to the maintenance of this functional resilience.

## DISCUSSION

We used the natural genetic variation of five closely related species to investigate and characterize features of CTCF binding at TAD boundaries. Our analyses reveal that CTCF binding sites at the boundaries of TADs are generally subject to stronger sequence constraints compared to CTCF sites in the background genome. Nevertheless, the CTCF binding profile at TAD borders seems to also be evolving under the effect of dynamic evolutionary processes. This is indicated by numerous gains of new species-specific CTCF binding sites close to species-conserved ones, giving rise to mixed clusters containing both evolutionary old and young CTCF binding sites.

Our data show that CTCF binding is largely conserved across *Mus* species, consistent with prior studies that demonstrate conservation across mammals (Kunarso et al., 2010; Schmidt et al., 2010, 2012). Our data also indicate that the boundaries of TADs commonly overlap with *Mus*-conserved CTCF sites, similar to observations from more distantly related mammalian lineages (Rao et al., 2014; Vietri Rudan et al., 2015). We show that a significant fraction of species-specific CTCF sites also localize in the vicinity of TAD borders, and that CTCF binding sites at TAD boundaries have both stronger sequence constraints and stronger binding affinity, independent of their conservation across species. Our data also reveal discrepancies in the expansion of TE classes at TAD boundary regions compared to the background genome. Specifically, TAD boundaries are relatively depleted of both LINE elements and LINE-derived CTCF binding sites, suggesting negative selection against insertions of long—and potentially disrupting—sequences at TAD boundaries. This is complementary to observed structural variant depletion at TAD boundaries as an effect of purifying selection (Fudenberg and Pollard, 2019). Overall, these observations suggest that the functional role of CTCF binding at TAD boundary regions is maintained by multiple evolutionary mechanisms including local sequence constraint, new site acquisition, and rejection of insertions and deletions.

Our results show that dynamically conserved regions that contain clusters of CTCF sites are another common characteristic of TAD boundaries. These clusters comprise both conserved CTCF binding events, which were apparently fixed at TAD boundary regions in the common ancestor, and divergent sites, which are the result of more recent gains or losses within the distinct mouse lineages. These clusters suggest a mechanism by which local turnover events can largely preserve TAD structure and function. Indeed, a recent study has demonstrated CTCF binding site turnover at loop anchors mediated by TEs, and it suggested that this is a common mechanism of contributing to conserved genome folding events between human and mouse (Choudhary et al., 2018). Based on these observations, we conclude that the formation of CTCF binding site clusters serves as an additional evolutionary buffering mechanism to preserve the CTCF binding potential of TAD boundaries and ensure resilience of higher order chromatin structure by maintaining a dynamic redundancy of CTCF binding sites.

Evolutionarily conserved clusters of CTCF binding sites may help explain previous observations of TAD structures remaining intact upon experimental disruption of individual or multiple CTCF sites, assuming that such clustered CTCF binding sites can be used interchangeably to provide higher order resilience against local disruptions. For example, Nora et al. showed that the deletion of a TAD boundary is followed by ectopic *cis*-interactions locally but adjacent TADs do not merge; they hypothesize that there must be additional elements within TADs that “act as relays when a main boundary is removed” (Nora et al., 2012). Furthermore, Barutcu et al. demonstrated that TAD structures are preserved upon deletion of the CTCF-rich *Firre* locus from a TAD boundary (Barutcu et al., 2018). They hypothesize that additional CTCF binding sites outside the *Firre* locus may serve to recruit CTCF and thus help maintain the TAD boundary. In addition, a recent study on CTCF hemizygosity suggested that, within genes, adjacent CTCF sites may have subtle additive effects on gene expression (Aitken et al., 2018), suggesting that clustered CTCF sites may enhance other CTCF functions. We also found that gene expression around TAD boundaries in cases of species-specific losses of individual CTCF sites is highly robust. As a whole, our results strongly suggest that the dynamic conservation of genomic regions harboring clusters of CTCF sites is an important feature of CTCF binding evolution, which is critical to the functional stability of higher order chromatin structure. Interestingly, such clusters are also found in genomic regions other than TAD borders. It is possible that these regions are related to the establishment of higher order chromatin structure, potentially representing unidentified TAD boundaries or loop anchors, or other functional and regulatory roles of CTCF.

Further insight into the functional implications of CTCF site clusters come from our result that CTCF clusters colocalize with the cohesin subunit RAD21 to a greater frequency than singleton CTCF sites. Moreover, we demonstrate that clustered CTCF sites are located significantly closer to TSSs than singleton sites. Together, these suggest that clusters play an important role in stabilizing cohesin at specific genomic regions, as well as in transcriptional regulation. These observations may provide new mechanistic insight to the previously proposed dynamic loop maintenance complex (LMC) model, in which cohesin associates with a genomic region for a significantly longer time than CTCF molecules (Hansen et al., 2017). Specifically, our observations of clustered CTCF binding sites support the proposed rapid unloading and rebinding of CTCF molecules in close genomic proximity, which facilitates rapid cohesin translocation on DNA between CTCF binding sites that act as occasionally permeable boundary elements (Davidson et al., 2016; Hansen et al., 2017). This process apparently facilitates gene transcription by allowing RNA polymerase II to push cohesin along gene bodies (Borrie et al., 2017; Davidson et al., 2016; Heinz et al., 2018).

Finally, it is tempting to speculate a connection between our identified clusters of closely located CTCF binding sites on the genome and the reportedly observed 3D “clusters” (or “hubs”) of CTCF protein molecules (Hansen et al., 2018b, 2018c). In particular, Hansen et al. have proposed a guided mechanism where an RNA strand can bind to and gather together multiple CTCF protein molecules near cognate binding sites. These CTCF molecule hubs apparently enhance the search for target binding sites, increase the binding rate of CTCF to its related sites (also as part of the LMC model) and are often implicated in chromatin loop formation (Hansen et al., 2018b, 2018c). It is possible that our identified CTCF site clusters act synergistically with this mechanism as nearby sites for the concentrated CTCF molecules to bind.

In conclusion, we identified dynamic evolutionary clusters of CTCF binding sites as a feature of TAD boundary architecture and we propose that these likely contribute to the remarkable resilience of TAD structures and gene expression to losses and gains of individual CTCF binding sites. Thus, further studies of seeking a definitive understanding of the functional roles of CTCF might require consideration of extended regions that harbor clusters of multiple CTCF sites.

## METHODS

### ChIP-seq experiments and data analysis

To characterize the CTCF binding profile in *Mus musculus castaneus* (CAST/EiJ) and *M. spretus* (SPRET/EiJ), we performed chromatin immunoprecipitation experiments followed by high-throughput sequencing (ChIP-seq) using adult liver tissue. ChIP-seq libraries and input control libraries from three biological replicates of each species were prepared as described in Schmidt et al., 2009. Subsequently, libraries were sequenced on a HiSeq2000 (Illumina) to produce 100bp paired-end sequence fragments.

In addition, we obtained published CTCF ChIP-seq data from the livers of *Mus musculus domesticus* (C57BL/6J), *Mus caroli*/EiJ, and *M. pahari*/EiJ (Thybert et al., 2018). Three biological replicates from each species were used.

We aligned sequenced reads from CAST and *M. spretus* to the reference genome assemblies CAST_EiJ_v1 and SPRET_EiJ_v1 (Lilue et al., 2018), respectively, with BWA mem version 0.7.12 (Li and Durbin, 2010) discarding reads with more than three occurrences. We also mapped the retrieved raw ChIP-seq reads from C57BL/6J, *M. caroli* and *M. pahari* to the genomes GRCm38 (mm10), CAROLI_EIJ_v1.1, and PAHARI_EIJ_v1.1 (Cunningham et al., 2019; Lilue et al., 2018), respectively, using the same method for the sake of performing matched analyses in all species. CTCF enrichment peaks were called with MACS 1.4.2 (Zhang et al., 2008) with a p-value threshold of 0.001. For downstream analyses, we used peaks identified in at least two replicates of each species.

We also performed ChIP-seq in C57BL/6J liver to identify genomic regions enriched for the cohesin subunit RAD21, using also an input control library from C57BL/6J liver from Thybert et al., 2018. Sample preparation and chromatin immunoprecipitation was performed as described in Schmidt et al., 2012 using 10μg RAD21 antibody (Abcam, ab992, lot GR12688-8). Immunoprecipitated DNA and 50ng of input DNA was used for library preparation using the ThruPLEX DNA-Seq library preparation protocol (Rubicon Genomics, UK). Library fragment size was determined using a 2100 Bioanalyzer (Agilent). Libraries were quantified by qPCR (Kapa Biosystems). Pooled libraries were deeply sequenced on a HiSeq2500 (Illumina) according to manufacturer’s instructions to produce single-end 50bp reads. We obtained sequenced reads and mapped them to the mouse genome assembly GRCm38 using BWA 0.6.1 (Li and Durbin, 2010). We then called RAD21 peaks using MACS2 2.1.2.1 with default options (Zhang et al., 2008).

### TADs

We used the boundaries of mouse liver TADs published by Vietri Rudan et al., 2015. We considered as TAD boundaries the start and end nucleotides of each TAD, while in some of the analyses (where indicated in the following methods description) we used a window of +/-50kb around them to study TAD boundary regions.

### Conservation of CTCF binding sites in *Mus* species

To investigate the conservation of CTCF binding across the studied *Mus* species, we first found the orthologous alignments of the CTCF ChIP-seq peaks in the genomes of the other species. These orthologous CTCF regions across mice were obtained using an extended version of the eutherian mammal Endo-Pecan-Ortheus (EPO) multiple genome alignment that also included the genomes of CAST, *M. spretus, M.caroli, and M. pahari* (Thybert et al., 2018). Once the orthologous regions of CTCF sites were identified in all *Mus* species, we cross-validated the binding of CTCF in each species using the corresponding ChIP-seq data. Specifically, we considered that a CTCF site was conserved if it (a) it had an orthologous alignment across species and (b) the orthologous alignments also contained a CTCF ChIP-seq peak (Fig. 1B).

### Binding affinity and sequence constraint of CTCF motifs

To identify CTCF binding motifs, we retrieved the FASTA sequences of all CTCF peaks in C57BL/6J, using bedtools getfasta v.2.25.0 (Quinlan and Hall, 2010) and scanned these sequences for the primary CTCF binding motif (M1) from the JASPAR database (Mathelier et al., 2014) using Find Individual Motif Occurrences (FIMO) from the MEME suite v.4.12.0 (Bailey et al., 2009; Grant et al., 2011) with default parameters. We extended the identified 19 base-long M1 motifs to include 20 bases upstream and 20 bases downstream in order to allow discovery of the extended version of the motifs (M1 and M2). Finally, we calculated the binding affinity of these sequences for CTCF using DeepBind v.0.11 (Alipanahi et al., 2015), as in Aitken et al., 2018, and compared the significance of the difference between distributions of the affinity values between motifs found in TAD-boundary-associated and non-TAD-boundary-associated CTCF peaks at each conservation level (Fig. 2A-B).

To retrieve Rejected Substitution (RS) scores for each position of every identified 19 base-long M1 motif in C57BL/6J, we obtained pre-calculated GERP (Cooper, 2005) conservation scores for each nucleotide of these mouse M1 sequences from Ensembl (Herrero et al., 2016). The RS score of a genomic position was calculated as the difference of observed to expected substitutions. We then averaged the RS score per position among all motifs and compared these averaged RS scores of TAD-boundary-associated M1 motifs with non-TAD-boundary-associated motifs (Fig. 2E, 2F).

### ChIP-seq enrichment of identified CTCF peaks

The CTCF sites that we identified in each species were the intersection of the CTCF peaks called in ≥ 2 biological replicates. We calculated the ChIP-seq fragment enrichment of each CTCF site by averaging the ChIP enrichment scores, reported by MACS, over the replicates. We then compared the significance of the difference between the distributions of average ChIP enrichment between TAD-boundary-associated and non-TAD-boundary-associated CTCF sites of each conservation level using Mann-Whitney U tests (Fig. 2C, 2D).

### Motif word usage analysis

We scanned all CTCF peaks from each of the five species for the primary CTCF binding motif (M1) using FIMO from the MEME suite as described above. From the 19-base M1 motif instances identified in each species we retrieved the central most informative 14-mer and estimated its frequency of occurrence as: the number of occurrences of the 14-mer word in CTCF binding regions divided by the number of occurrences of the word in the whole genome of the species using the procedure of Schmidt et al., 2012. We filtered out any motif word that occurred fewer than five times in the whole genome. We illustrated the occurrence frequency of the motif words in each species on a heatmap which is sorted by distance to the closest TAD border (Fig. S4).

### Association of CTCF binding sites with classes of transposable elements

We used the full set of CTCF sites identified in all species and projected them on to the C57BL/6J genome (GRCm38), as well as published transposable elements in C57BL/6J (Thybert et al., 2018; https://www.ebi.ac.uk/research/flicek/publications/FOG21). We intersected the center of each CTCF binding site with the transposable elements and reported the number of CTCF site centers that overlapped with each TE class. The overall representation of each TE class in the whole genome that is shown as a reference (marked as “background” in Fig. 3A) was calculated as: the total length of all TEs belonging to each class (SINE, LINE, LTR, DNA) sequences divided by the total genome length.

### Representation of TE classes at TAD boundary regions

As for Fig. 3B, we defined TAD boundary regions as genomic windows of 50kb upstream and 50kb downstream of the boundaries of TADs. To evaluate the representation of each TE class, we summed the length of sequences corresponding to each TE class that occurred within each TAD boundary region and divided that by the total length of the TAD boundary region, i.e.100kb. To retrieve random genomic regions of similar length and distribution, we shuffled the TAD boundary regions using bedtools shuffle v2.2.5.0, having first excluded chromosome Y, genome scaffolds, and chromosome ends, where TADs are not called. We repeated the same calculation for TE class representation as above for these shuffled TAD boundaries, i.e. random genomic regions. We then plotted the distribution of these values for TAD boundary regions and random genomic regions. To determine the representation of each TE class in the background genome (dotted line in Fig. 3B), we divided again the total length of all sequences that correspond to each TE class by the total C57BL/6J genome (GRCm38) length, analogous to the CTCF TE class analysis above.

### Density of CTCF sites at TAD boundaries and clusters of CTCF binding sites

To determine the enrichment of CTCF binding sites in TAD boundary regions (compared to the surrounding genome) we measured the distance of each CTCF binding site to its closest TAD boundary using bedtools closest. We then categorized the CTCF sites based on their conservation level. For each CTCF site conservation level, we grouped all distance values up to +/-300kb in bins of 20kb and plotted the number of CTCF sites in each bin divided by the length of the bin, i.e. 20kb (Fig. 4A). To further characterize the density of CTCF sites at TAD boundaries, we grouped CTCF sites both according to their conservation level and association with a TAD boundary (vs. no association with any TAD boundary), and for each of these categories we found the distance of each CTCF site from its closest CTCF site using bedtools closest (Fig. 4B).

To identify clusters of CTCF binding sites, we used the full set of CTCF binding sites of all five *Mus* species projected onto the C57BL/6J genome (GRCm38/mm10), as shown in Fig. 1B. We identified instances of consecutive CTCF sites that were up to 10kb apart from each other, using bedtools cluster. We then determined and compared the enrichment of clustered and singleton CTCF sites at TAD boundaries using the same approach as in Fig. 4A but having categorized the CTCF sites based on whether they belong to a cluster (clustered) or not (singletons) (Fig. 4C).

For Figures 4D and 4E, we again defined TAD boundary regions as TAD boundary +/-50kb. We categorized these regions based on the *highest* conservation level of their CTCF sites. Subsequently, for each category we counted its total number of CTCF sites (Fig. 4D), as well as the number of these TAD boundary regions with clustered CTCF sites and with only singleton sites (Fig. 4E).

For Fig. S5, we defined *Mus-*conserved (5-way) CTCF sites with a distance to the closest TAD border >80kb as non-TAD-boundary associated. We calculated the enrichment of 1-way (species-specific), 2-way, 3-way, and 4-way conserved CTCF sites in their vicinity in the same way as in for TAD boundaries (Fig. 4A), but using as anchor the non-TAD-boundary-associated 5-way CTCF sites themselves, instead of the TAD boundaries.

### Clusters in C57BL/6J and cluster conservation analyses

We identified clusters of CTCF binding sites in C57BL/6J (Fig. S6) in the same way as for Fig. 4C but using only CTCF peaks called in C57BL/6J. We used the same methods as for Fig. 4A and 4C to determine the enrichment of CTCF sites of different conservation levels at TAD borders (Fig. S6A), as well as the enrichment of clustered versus singleton CTCF sites (Fig. S6B).

To estimate the conservation of CTCF sites clusters (Fig. S6D), we identified all the genomic regions that correspond to clusters of CTCF sites in each of the five species separately. We then projected through whole-genome alignments (see “Conservation of CTCF binding sites in Mus species” Methods section) the cluster regions of each species onto the C57BL/6J genome and determined whether they overlap with the orthologous cluster regions of the other species.

### RNA-seq data

We retrieved published liver-derived RNA-seq data from six biological replicates for each of the species C57BL/6J and *M. m. castaneus* (Goncalves et al., 2012), as well as from four biological replicates of *M. caroli* (Wong et al., 2015). To have the same number of replicates in each spcies, we further generated and sequenced two additional RNA-seq libraries for *M. caroli* following the methods described in Goncalves et al., 2012 and Wong et al., 2015. Briefly, total RNA was extracted from two independent liver samples using Qiazol (Qiagen) and DNase treated with DNA-free DNA Removal Kit (Ambion). Polyadenylated mRNA was enriched, directional double-stranded cDNA was generated, fragmented by sonication, and prepared for sequencing. Each of the two libraries were sequenced on an Illumina GAIIx to generate 75bp paired-end fragments.

### RNA-seq data processing and analysis

Adapter sequences were trimmed off with reaper from the Kraken tool suite (Davis et al., 2013). The paired-end RNA-seq reads from each replicate of C57BL/6J, CAST, and *M. caroli* were mapped to the corresponding species genomes (see “ChIP-seq experiments and data analysis” Methods section) using STAR 1.5.2 (Dobin et al., 2013) with default settings. Raw reads mapping to annotated genes were counted using htseq-count (Anders et al., 2015). We then used the raw read counts to perform differential expression analyses with DESeq2 1.20.0 (Love et al., 2014) with default settings.

To determine gene expression patterns around instances of 5-way conserved CTCF sites and species-specific CTCF site losses at TAD boundaries (Fig. 6A, 6D and 6G), we first identified the closest upstream and downstream gene in each species using the gene annotation from Ensembl version 95 (Cunningham et al., 2019) and then calculated the relative gene expression of downstream to upstream gene in each species. We were not interested in the relative expression of the gene pair flanking a CTCF site *per se*, but in whether this ratio for each CTCF site is consistent between species when the in-between CTCF binding separating them changes. For this reason, we only used CTCF sites that were flanked by 1:1 orthologous genes between the three species. We went on to use DESeq2 (Love et al., 2014) in order to compute the log_2_(fold change) between the downstream and upstream gene – as a measure of the relative expression of genes flanking each CTCF site – in each species, and to subsequently compare this log_2_(fold change) between species. Since DESeq2 is not designed to normalize for gene lengths, and our aim was to generate comparable expression pattern estimations between the species, we also required all the orthologous genes that we used to have a similar length among the three species (0.7 < *len_ratio < 1.3,* where *len_ratio* is the length of gene in species A divided by the length of its orthologous gene in species B). Finally, we compared the calculated log2(fold-change) values for each gene pair in C57BL/6J with the corresponding value of its orthologous gene pair in CAST (Fig. 6B, 6E, 6H) and in *M. caroli* (Fig. 6C, 6F, 6I).

## DATA ACCESS

All ChIP-seq and RNA-seq data generated in this study have been submitted to Array Express (https://www.ebi.ac.uk/arrayexpress/) under accession numbers E-MTAB-8014 and E-MTAB-8016, respectively.

## AUTHOR CONTRIBUTIONS

EK, DTO, MR, and PF conceived and designed the study. EK led and conducted data analysis with contributions from MR. SJA, CF, and KS generated data. SJA and XI-S generated and shared additional unpublished data and analysis. EK, SJA, DTO, MR, and PF wrote the manuscript and created figures. All authors read the final manuscript and provided critical comments.

## ACKNOWLEDGMENTS

Funding provided by Wellcome Trust (WT108749/Z/15/Z, WT202878/Z/16/Z, WT202878/B/16/Z, and WT106563/Z/14 to SJA.), Cancer Research UK (20412), the European Research Council (615584), the European Molecular Biology Laboratory, and the EMBL International PhD Programme. We thank John Marioni, Vasavi Sundaram, Margus Lukk, and Matthew P. Davis for support and helpful discussions.

PF is a member of the Scientific Advisory Boards of Fabric Genomics, Inc., and Eagle Genomics, Ltd.

## SUPPLEMENTARY FIGURES

**Figure S1:**
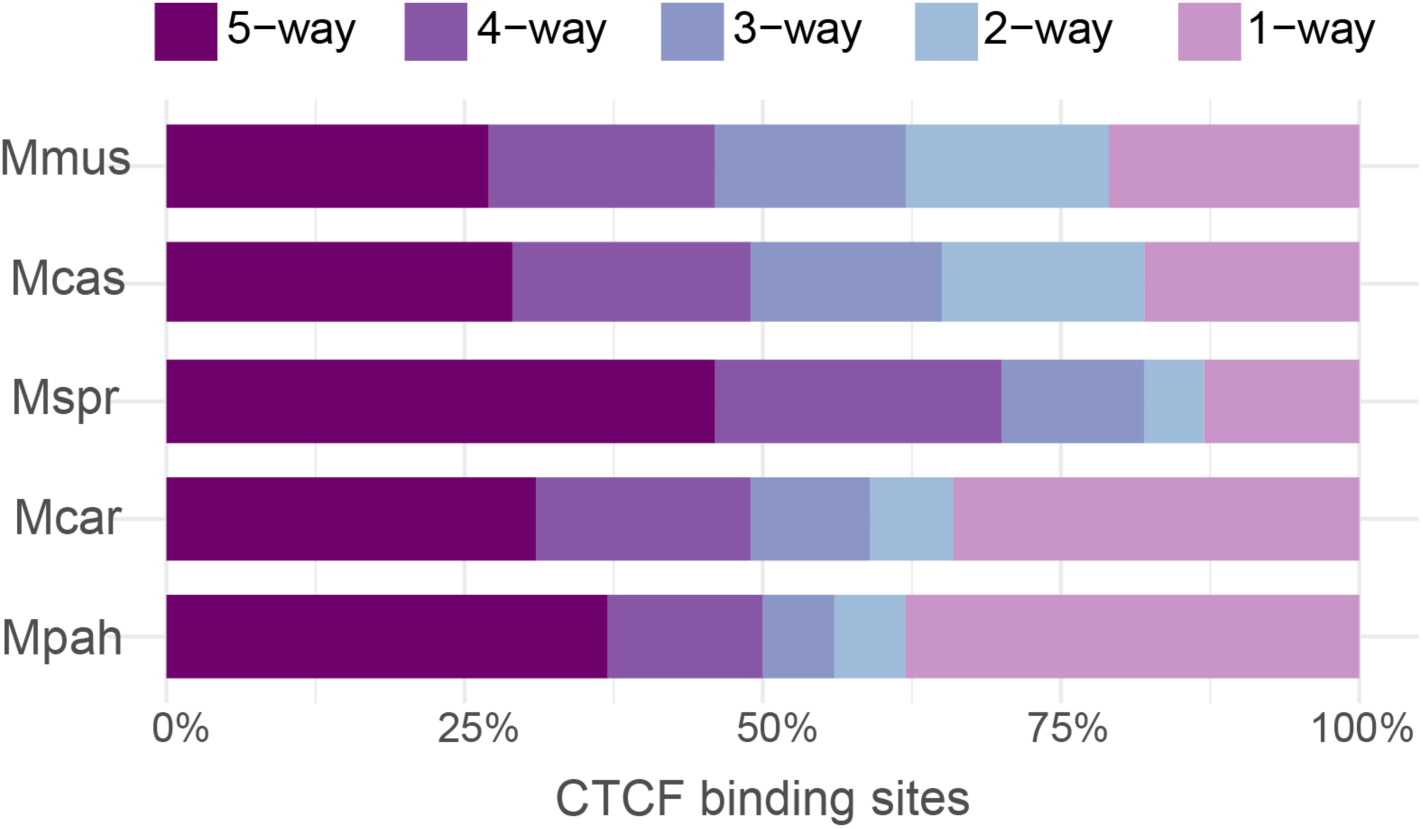
Fractions of CTCF binding sites of different conservation levels in each of the studied *Mus* species.

**Figure S2:**
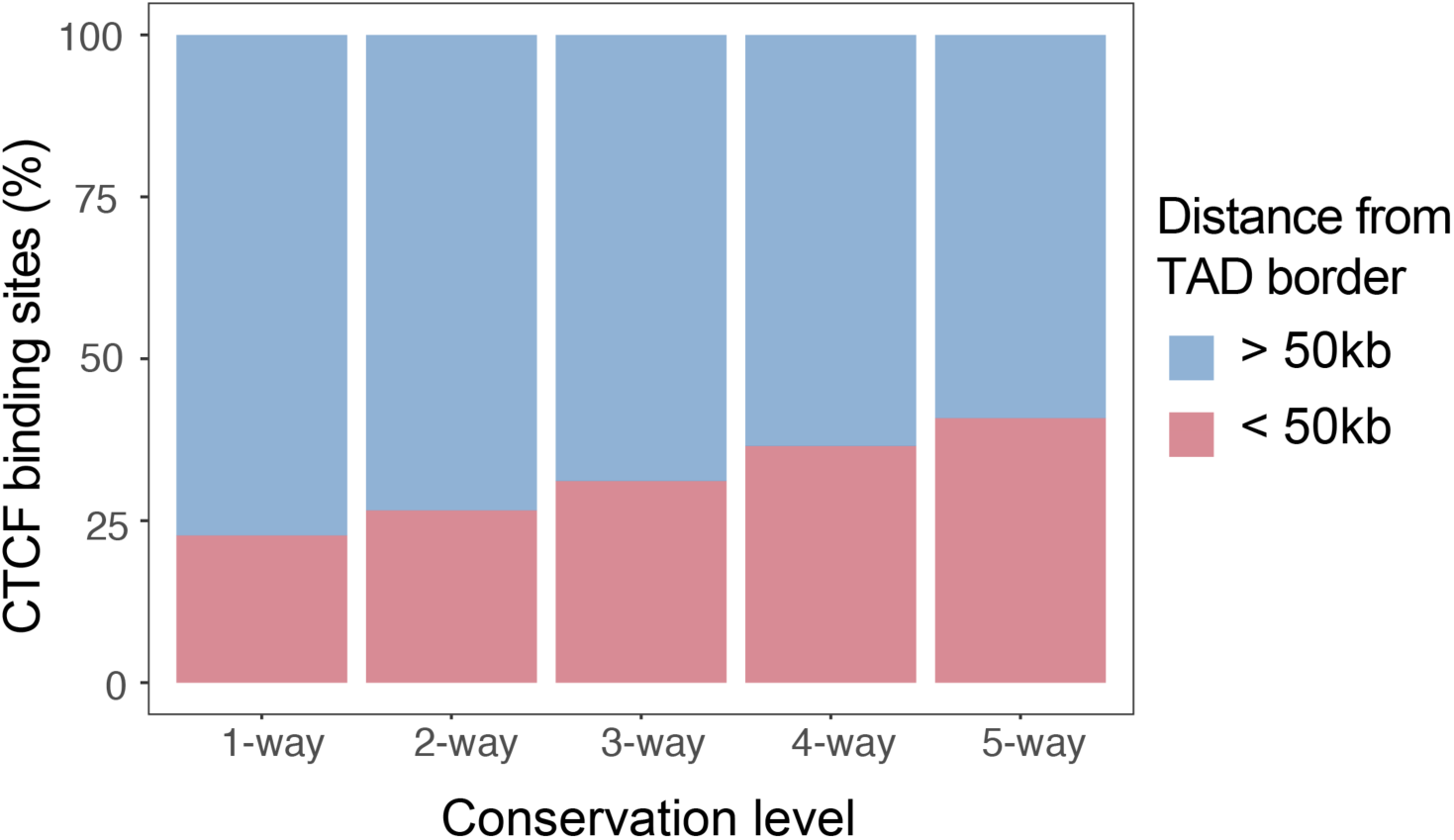
Fractions of all *Mus* CTCF sites of each conservation level that are associated (*d*≤ 50kb) or not associated (*d* > 50kb) with TAD boundaries.

**Figure S3:**
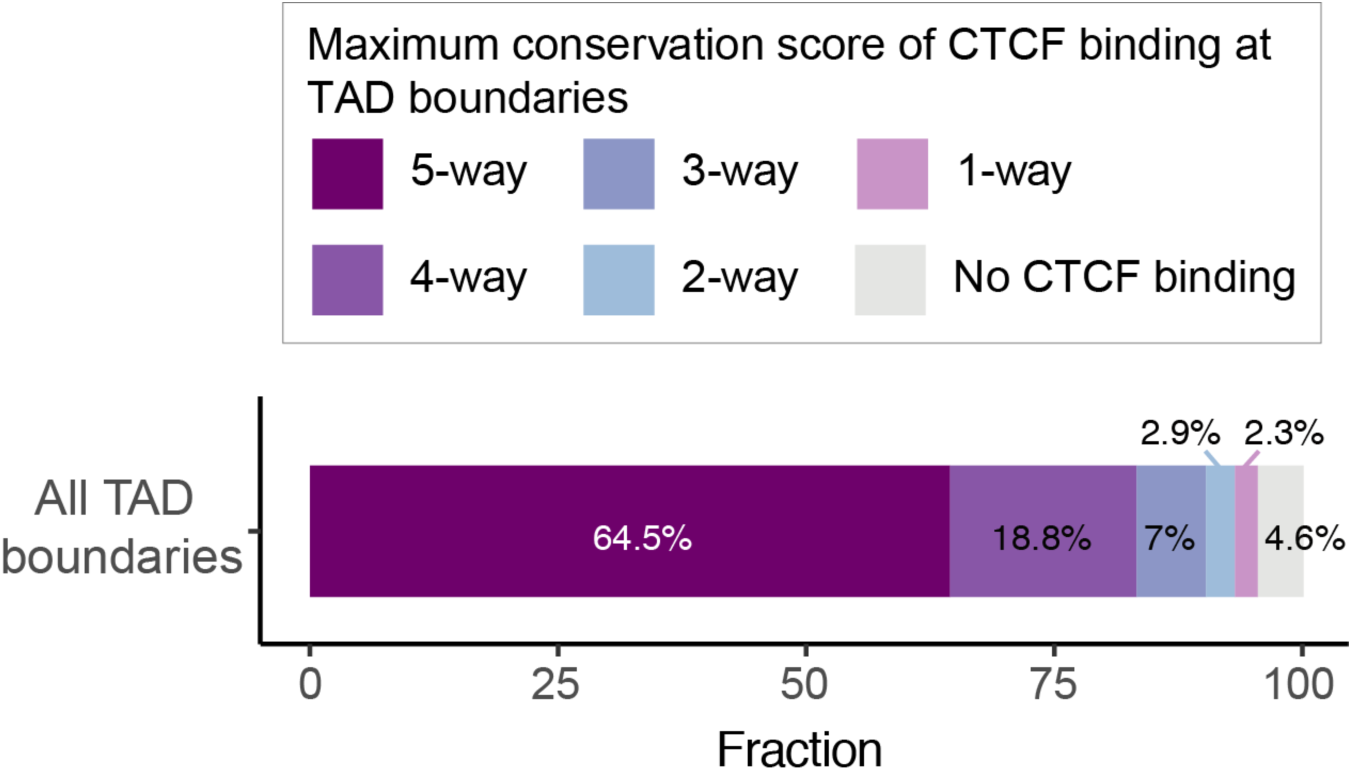
Fractions of TAD boundaries with CTCF sites of different conservation levels. Most TAD boundaries (64%) harbor at least one *Mus*-conserved (5-way) CTCF site. Lower percentages of TAD borders do not contain any *Mus-*conserved CTCF site but overlap with less conserved sites or do not bind CTCF at all.

**Figure S4:**
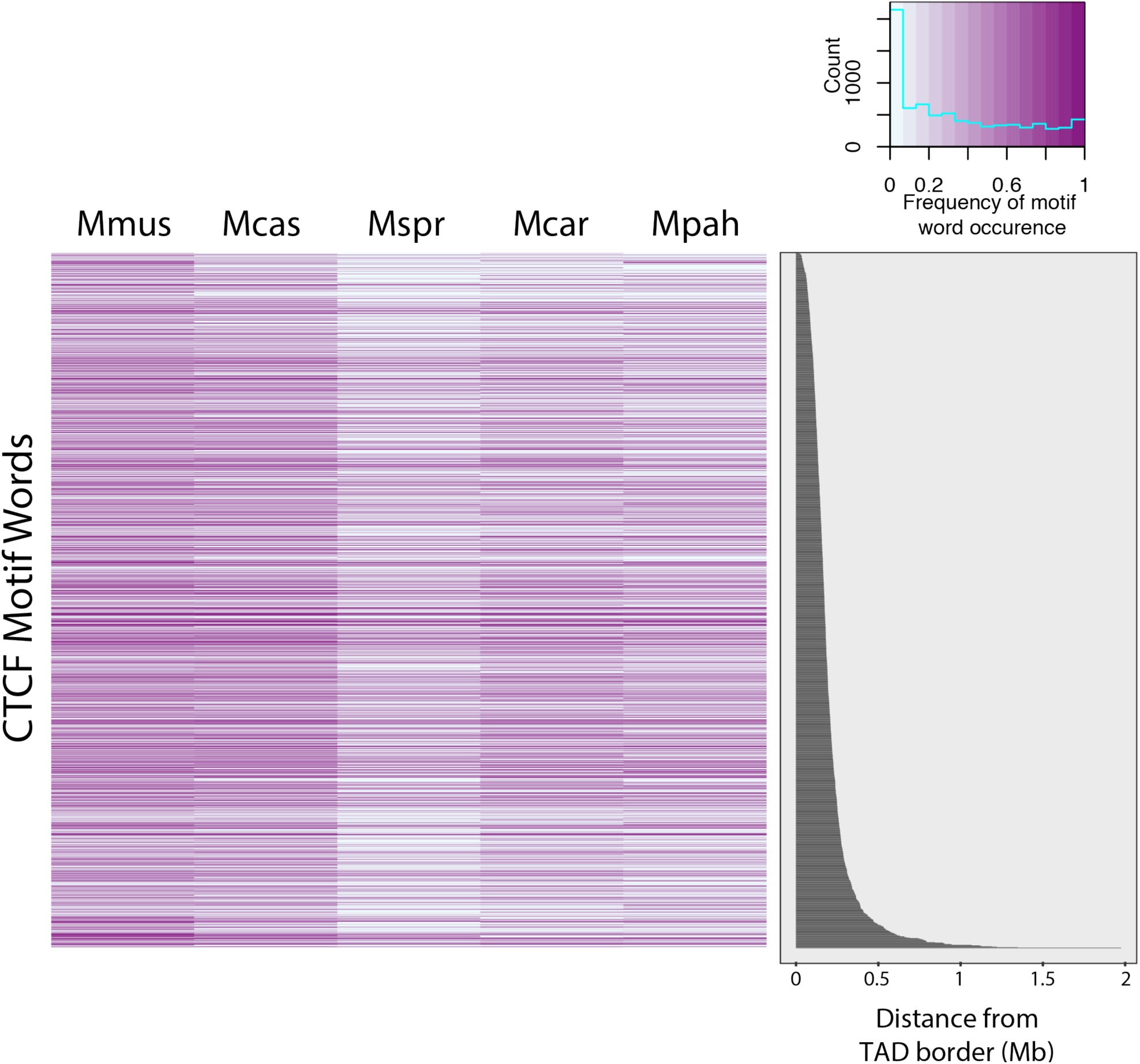
There is no evidence of any enrichment of specific motif words at TAD boundaries among the species. Heatmap of the ∼1,500 motif words found in CTCF peaks in the five *Mus* species. Each row corresponds to a motif word, while the color density represents its frequency of occurrence. The occurrence frequency of each motif word in the CTCF peaks is normalized by the number of its occurrences in the whole genome for the respective species. Motif words in the heatmap are sorted based on their distance to the closest TAD boundary. There is no evidence of any selected set of motif words being used with significant frequency at TAD boundaries among the species. The lower density of motif words is *M. spretus* reflects the smaller number of CTCF binding sites identified in that species.

**Figure S5:**
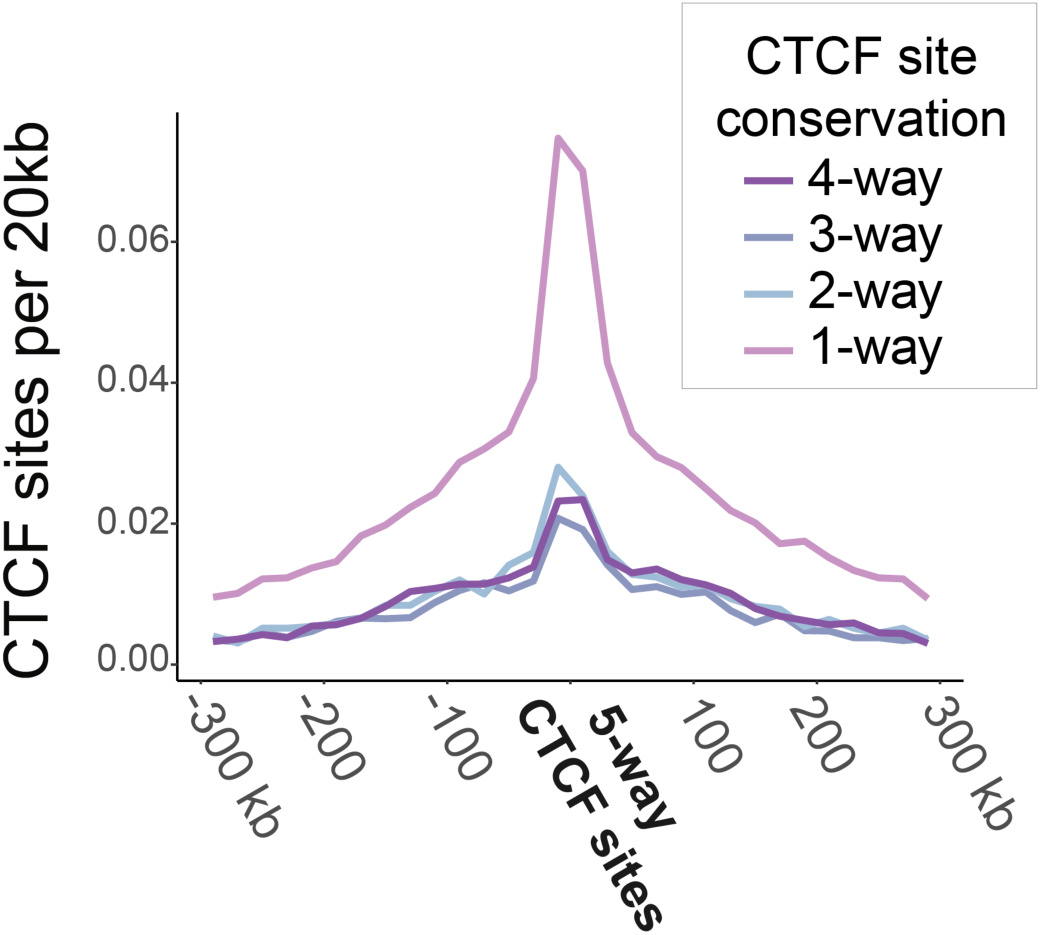
Clusters of both conserved and species-specific CTCF sites might also occur away from TAD boundaries. Enrichment of CTCF sites of different conservation levels around *Mus*-conserved CTCF sites that are *not* associated with TAD boundaries (distance from closest TAD border: *d* > 80kb). A high number of species-specific (1-way) CTCF sites are concentrated around these “anchor” 5-way conserved sites, showing that sites of mixed conservation levels can be clustered together.

**Figure S6:**
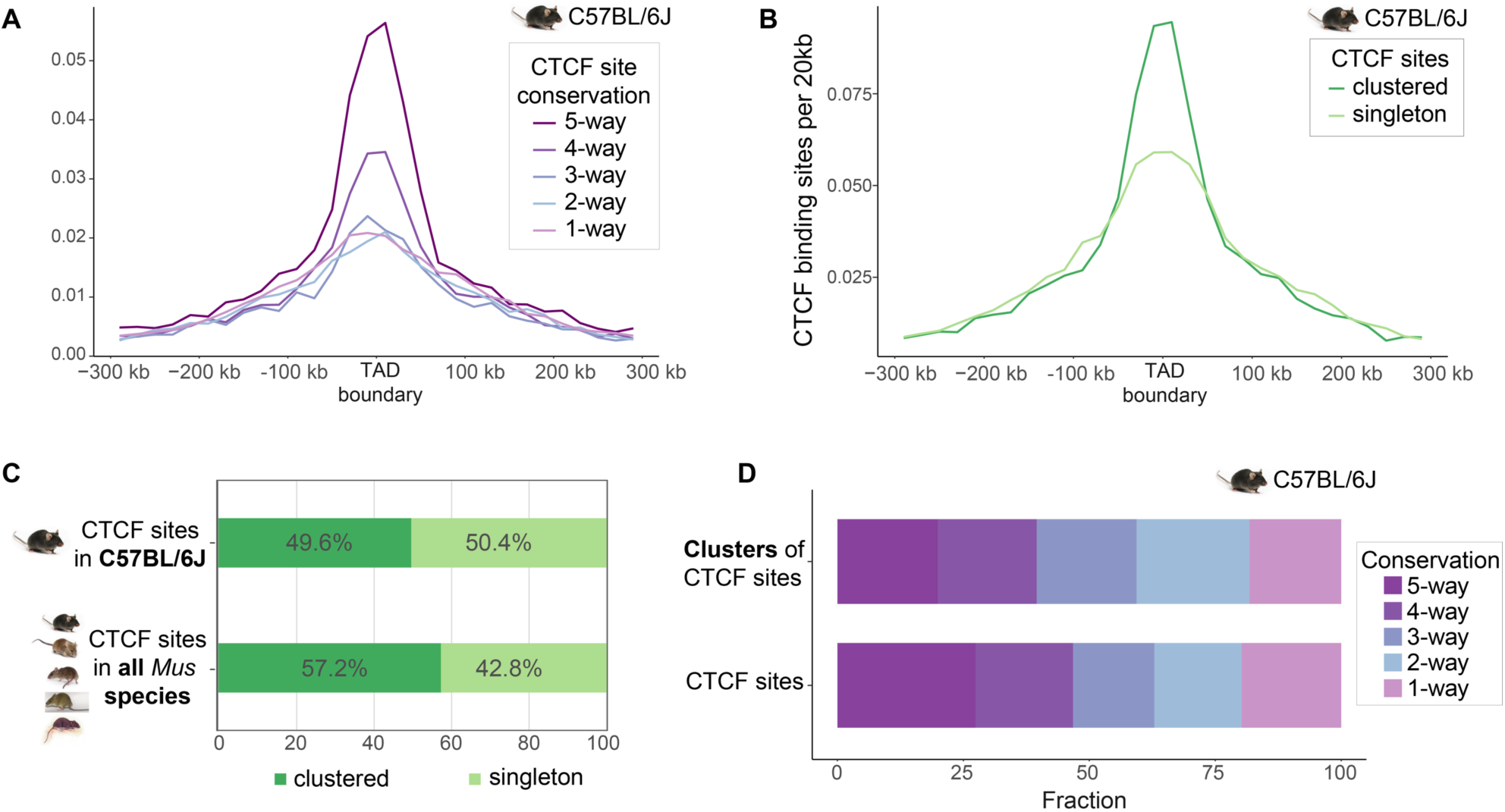
Inspection of the CTCF binding profile in C57BL/6J confirms that CTCF sites form clusters in individual species. (A) Enrichment of C57BL/6J CTCF sites of different conservation levels at TAD boundaries. (B) Clustered C57BL/6J CTCF sites are more highly enriched than singleton sites at TAD borders. (C) The fraction of clustered CTCF sites in C57BL/6J is similar to that of CTCF sites belonging to ancestral *Mus* clusters. (D) The conservation pattern of CTCF site clusters, as distinct functional entities, resembles that of individual CTCF binding sites.

**Figure S7:**
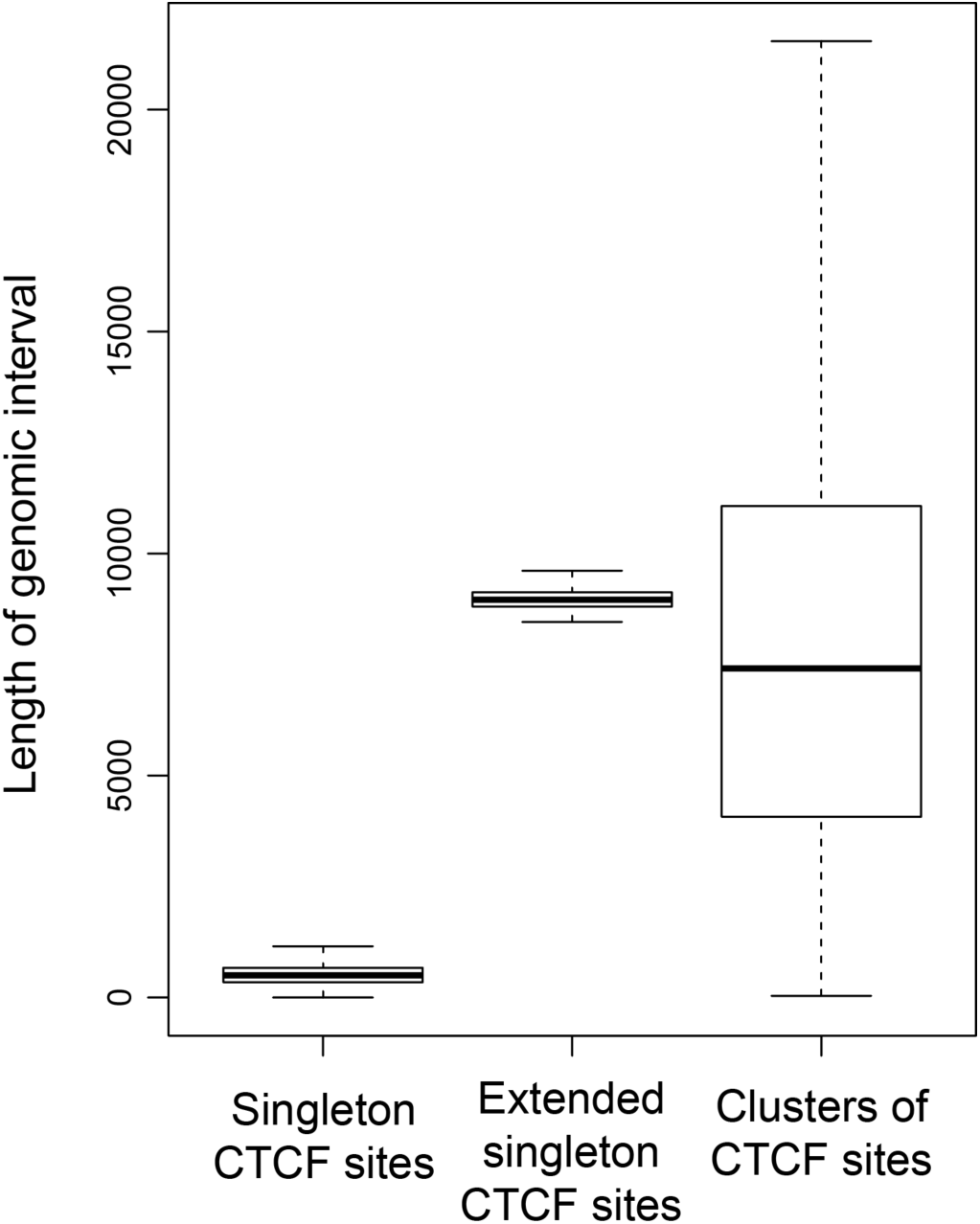
Length distribution of genomic intervals occupied by singleton CTCF sites, “extended” singleton CTCF sites and clusters of CTCF sites. The extended singleton CTCF sites represent genomic windows of singleton CTCF sites that were extended so that the mean of their length distribution becomes equal to that of the length distribution for the CTCF clusters.

## REFERENCES

Aitken, S.J., Ibarra-Soria, X., Kentepozidou, E., Flicek, P., Feig, C., Marioni, J.C., and Odom, D.T. (2018). CTCF maintains regulatory homeostasis of cancer pathways. Genome Biol. 19, 106.

Alipanahi, B., Delong, A., Weirauch, M.T., and Frey, B.J. (2015). Predicting the sequence specificities of DNA- and RNA-binding proteins by deep learning. Nat. Biotechnol. 33, 831–838.

Anders, S., Pyl, P.T., and Huber, W. (2015). HTSeq--a Python framework to work with high-throughput sequencing data. Bioinformatics 31, 166–169.

Bailey, T.L., Boden, M., Buske, F.A., Frith, M., Grant, C.E., Clementi, L., Ren, J., Li, W.W., and Noble, W.S. (2009). MEME SUITE: tools for motif discovery and searching. Nucleic Acids Res. 37, W202–W208.

Baniahmad, A., Steiner, C., Köhne, A.C., and Renkawitz, R. (1990). Modular structure of a chicken lysozyme silencer: Involvement of an unusual thyroid hormone receptor binding site. Cell 61, 505–514.

Barutcu, A.R., Maass, P.G., Lewandowski, J.P., Weiner, C.L., and Rinn, J.L. (2018). A TAD boundary is preserved upon deletion of the CTCF-rich Firre locus. Nat. Commun. 9, 1444.

Borrie, M.S., Campor, J.S., Joshi, H., and Gartenberg, M.R. (2017). Binding, sliding, and function of cohesin during transcriptional activation. Proc. Natl. Acad. Sci. 114, E1062–E1071.

Bourque, G., Leong, B., Vega, V.B., Chen, X., Lee, Y.L., Srinivasan, K.G., Chew, J.-L., Ruan, Y., Wei, C.-L., Ng, H.H., et al. (2008). Evolution of the mammalian transcription factor binding repertoire via transposable elements. Genome Res. 18, 1752–1762.

Chen, H., Tian, Y., Shu, W., Bo, X., and Wang, S. (2012). Comprehensive Identification and Annotation of Cell Type-Specific and Ubiquitous CTCF-Binding Sites in the Human Genome. PLoS One 7, e41374.

Choudhary, M.N., Friedman, R.Z., Wang, J.T., Jang, H.S., Zhuo, X., and Wang, T. (2018). Co-opted transposons help perpetuate conserved higher-order chromosomal structures. BioRxiv 485342.

Cooper, G.M. (2005). Distribution and intensity of constraint in mammalian genomic sequence. Genome Res. 15, 901–913.

Cunningham, F., Achuthan, P., Akanni, W., Allen, J., Amode, M.R., Armean, I.M., Bennett, R., Bhai, J., Billis, K., Boddu, S., et al. (2019). Ensembl 2019. Nucleic Acids Res. 47, D745–D751.

Davidson, I.F., Goetz, D., Zaczek, M.P., Molodtsov, M.I., Huis in’t Veld, P.J., Weissmann, F., Litos, G., Cisneros, D.A., Ocampo-Hafalla, M., Ladurner, R., et al. (2016). Rapid movement and transcriptional re-localization of human cohesin on DNA. EMBO J. 35, 2671–2685.

Davis, M.P.A., van Dongen, S., Abreu-Goodger, C., Bartonicek, N., and Enright, A.J. (2013). Kraken: A set of tools for quality control and analysis of high-throughput sequence data. Methods 63, 41–49.

Dixon, J.R., Selvaraj, S., Yue, F., Kim, A., Li, Y., Shen, Y., Hu, M., Liu, J.S., and Ren, B. (2012). Topological domains in mammalian genomes identified by analysis of chromatin interactions. Nature 485, 376–380.

Dobin, A., Davis, C.A., Schlesinger, F., Drenkow, J., Zaleski, C., Jha, S., Batut, P., Chaisson, M., and Gingeras, T.R. (2013). STAR: ultrafast universal RNA-seq aligner. Bioinformatics 29, 15–21.

Filippova, G.N., Fagerlie, S., Klenova, E.M., Myers, C., Dehner, Y., Goodwin, G., Neiman, P.E., Collins, S.J., and Lobanenkov, V. V (1996). An exceptionally conserved transcriptional repressor, CTCF, employs different combinations of zinc fingers to bind diverged promoter sequences of avian and mammalian c-myc oncogenes. Mol. Cell. Biol. 16, 2802–2813.

Flavahan, W.A., Drier, Y., Liau, B.B., Gillespie, S.M., Venteicher, A.S., Stemmer-Rachamimov, A.O., Suvà, M.L., and Bernstein, B.E. (2016). Insulator dysfunction and oncogene activation in IDH mutant gliomas. Nature 529, 110–114.

Fudenberg, G., and Pollard, K.S. (2019). Chromatin features constrain structural variation across evolutionary timescales. Proc. Natl. Acad. Sci. 116, 2175–2180.

Fudenberg, G., Imakaev, M., Lu, C., Goloborodko, A., Abdennur, N., and Mirny, L.A. (2016). Formation of Chromosomal Domains by Loop Extrusion. Cell Rep. 15, 2038–2049.

Gasch, A.P., Payseur, B.A., and Pool, J.E. (2016). The Power of Natural Variation for Model Organism Biology. Trends Genet. 32, 147–154.

Gómez-Marín, C., Tena, J.J., Acemel, R.D., López-Mayorga, M., Naranjo, S., de la Calle-Mustienes, E., Maeso, I., Beccari, L., Aneas, I., Vielmas, E., et al. (2015). Evolutionary comparison reveals that diverging CTCF sites are signatures of ancestral topological associating domains borders. Proc. Natl. Acad. Sci. 112, 7542–7547.

Goncalves, A., Leigh-Brown, S., Thybert, D., Stefflova, K., Turro, E., Flicek, P., Brazma, A., Odom, D.T., and Marioni, J.C. (2012). Extensive compensatory cis-trans regulation in the evolution of mouse gene expression. Genome Res. 22, 2376–2384.

Grant, C.E., Bailey, T.L., and Noble, W.S. (2011). FIMO: scanning for occurrences of a given motif. Bioinformatics 27, 1017–1018.

Guo, Y., Xu, Q., Canzio, D., Shou, J., Li, J., Gorkin, D.U., Jung, I., Wu, H., Zhai, Y., Tang, Y., et al. (2015). CRISPR Inversion of CTCF Sites Alters Genome Topology and Enhancer/Promoter Function. Cell 162, 900–910.

Hansen, A.S., Pustova, I., Cattoglio, C., Tjian, R., and Darzacq, X. (2017). CTCF and cohesin regulate chromatin loop stability with distinct dynamics. Elife 6, e25776.

Hansen, A.S., Cattoglio, C., Darzacq, X., and Tjian, R. (2018a). Recent evidence that TADs and chromatin loops are dynamic structures. Nucleus 9, 20–32.

Hansen, A.S., Hsieh, T.-H.S., Cattoglio, C., Pustova, I., Darzacq, X., and Tjian, R. (2018b). An RNA-binding region regulates CTCF clustering and chromatin looping. BioRxiv 495432.

Hansen, A.S., Amitai, A., Cattoglio, C., Tjian, R., and Darzacq, X. (2018c). Guided nuclear exploration increases CTCF target search efficiency. BioRxiv 495457.

Heinz, S., Romanoski, C.E., Benner, C., Allison, K.A., Kaikkonen, M.U., Orozco, L.D., and Glass, C.K. (2013). Effect of natural genetic variation on enhancer selection and function. Nature 503, 487–492.

Heinz, S., Texari, L., Hayes, M.G.B., Urbanowski, M., Chang, M.W., Givarkes, N., Rialdi, A., White, K.M., Albrecht, R.A., Pache, L., et al. (2018). Transcription Elongation Can Affect Genome 3D Structure. Cell 174, 1522-1536.e22.

Herrero, J., Muffato, M., Beal, K., Fitzgerald, S., Gordon, L., Pignatelli, M., Vilella, A.J., Searle, S.M.J., Amode, R., Brent, S., et al. (2016). Ensembl comparative genomics resources. Database 2016, bav096.

Ibn-Salem, J., Köhler, S., Love, M.I., Chung, H.R., Huang, N., Hurles, M.E., Haendel, M., Washington, N.L., Smedley, D., Mungall, C.J., et al. (2014). Deletions of chromosomal regulatory boundaries are associated with congenital disease. Genome Biol. 15, 423.

Kemp, C.J., Moore, J.M., Moser, R., Bernard, B., Teater, M., Smith, L.E., Rabaia, N.A., Gurley, K.E., Guinney, J., Busch, S.E., et al. (2014). CTCF Haploinsufficiency Destabilizes DNA Methylation and Predisposes to Cancer. Cell Rep. 7, 1020–1029.

Klenova, E.M., Nicolas, R.H., Paterson, H.F., Carne, A.F., Heath, C.M., Goodwin, G.H., Neiman, P.E., and Lobanenkov, V. V (1993). CTCF, a conserved nuclear factor required for optimal transcriptional activity of the chicken c-myc gene, is an 11-Zn-finger protein differentially expressed in multiple forms. Mol. Cell. Biol. 13, 7612–7624.

Kubo, N., Ishii, H., Gorkin, D., Meitinger, F., Xiong, X., Fang, R., Liu, T., Ye, Z., Li, B., Dixon, J., et al. (2017). Preservation of Chromatin Organization after Acute Loss of CTCF in Mouse Embryonic Stem Cells. BioRxiv 118737.

Kunarso, G., Chia, N.-Y., Jeyakani, J., Hwang, C., Lu, X., Chan, Y.-S., Ng, H.-H., and Bourque, G. (2010). Transposable elements have rewired the core regulatory network of human embryonic stem cells. Nat. Genet. 42, 631–634.

Li, H., and Durbin, R. (2010). Fast and accurate long-read alignment with Burrows–Wheeler transform. Bioinformatics 26, 589–595.

Lilue, J., Doran, A.G., Fiddes, I.T., Abrudan, M., Armstrong, J., Bennett, R., Chow, W., Collins, J., Collins, S., Czechanski, A., et al. (2018). Sixteen diverse laboratory mouse reference genomes define strain-specific haplotypes and novel functional loci. Nat. Genet. 50, 1574–1583.

Lobanenkov, V. V, Nicolas, R.H., Adler, V. V, Paterson, H., Klenova, E.M., Polotskaja, A. V, and Goodwin, G.H. (1990). A novel sequence-specific DNA binding protein which interacts with three regularly spaced direct repeats of the CCCTC-motif in the 5’-flanking sequence of the chicken c-myc gene. Oncogene 5, 1743–1753.

Love, M.I., Huber, W., and Anders, S. (2014). Moderated estimation of fold change and dispersion for RNA-seq data with DESeq2. Genome Biol. 15, 550.

Lupiáñez, D.G., Kraft, K., Heinrich, V., Krawitz, P., Brancati, F., Klopocki, E., Horn, D., Kayserili, H., Opitz, J.M., Laxova, R., et al. (2015). Disruptions of topological chromatin domains cause pathogenic rewiring of gene-enhancer interactions. Cell 161, 1012–1025.

Lupiáñez, D.G., Spielmann, M., and Mundlos, S. (2016). Breaking TADs: How Alterations of Chromatin Domains Result in Disease. Trends Genet. 32, 225–237.

Mathelier, A., Zhao, X., Zhang, A.W., Parcy, F., Worsley-Hunt, R., Arenillas, D.J., Buchman, S., Chen, C.Y., Chou, A., Ienasescu, H., et al. (2014). JASPAR 2014: An extensively expanded and updated open-access database of transcription factor binding profiles. Nucleic Acids Res.

Merkenschlager, M., and Nora, E.P. (2016). CTCF and Cohesin in Genome Folding and Transcriptional Gene Regulation. Annu. Rev. Genomics Hum. Genet. 17, 17–43.

Mifsud, B., Tavares-Cadete, F., Young, A.N., Sugar, R., Schoenfelder, S., Ferreira, L., Wingett, S.W., Andrews, S., Grey, W., Ewels, P.A., et al. (2015). Mapping long-range promoter contacts in human cells with high-resolution capture Hi-C. Nat. Genet. 47, 598–606.

Moon, H., Filippova, G., Loukinov, D., Pugacheva, E., Chen, Q., Smith, S.T., Munhall, A., Grewe, B., Bartkuhn, M., Arnold, R., et al. (2005). CTCF is conserved from Drosophila to humans and confers enhancer blocking of the Fab-8 insulator. EMBO Rep. 6, 165–170.

Nora, E.P., Lajoie, B.R., Schulz, E.G., Giorgetti, L., Okamoto, I., Servant, N., Piolot, T., van Berkum, N.L., Meisig, J., Sedat, J., et al. (2012). Spatial partitioning of the regulatory landscape of the X-inactivation centre. Nature 485, 381–385.

Nora, E.P., Goloborodko, A., Valton, A.-L., Gibcus, J.H., Uebersohn, A., Abdennur, N., Dekker, J., Mirny, L.A., and Bruneau, B.G. (2017). Targeted Degradation of CTCF Decouples Local Insulation of Chromosome Domains from Genomic Compartmentalization. Cell 169, 930-944.e22.

Ohlsson, R., Renkawitz, R., and Lobanenkov, V. (2001). CTCF is a uniquely versatile transcription regulator linked to epigenetics and disease. Trends Genet. 17, 520–527.

Ong, C.-T., and Corces, V.G. (2014). CTCF: an architectural protein bridging genome topology and function. Nat. Rev. Genet. 15, 234–246.

Parelho, V., Hadjur, S., Spivakov, M., Leleu, M., Sauer, S., Gregson, H.C., Jarmuz, A., Canzonetta, C., Webster, Z., Nesterova, T., et al. (2008). Cohesins Functionally Associate with CTCF on Mammalian Chromosome Arms. Cell 132, 422–433.

Phillips-Cremins, J.E., Sauria, M.E.G., Sanyal, A., Gerasimova, T.I., Lajoie, B.R., Bell, J.S.K., Ong, C.-T., Hookway, T.A., Guo, C., Sun, Y., et al. (2013). Architectural protein subclasses shape 3D organization of genomes during lineage commitment. Cell 153, 1281–1295.

Pombo, A., and Dillon, N. (2015). Three-dimensional genome architecture: players and mechanisms. Nat. Rev. Mol. Cell Biol. 16, 245–257.

Quinlan, A.R., and Hall, I.M. (2010). BEDTools: a flexible suite of utilities for comparing genomic features. Bioinformatics 26, 841–842.

Rao, S.S.P., Huntley, M.H., Durand, N.C., Stamenova, E.K., Bochkov, I.D., Robinson, J.T., Sanborn, A.L., Machol, I., Omer, A.D., Lander, E.S., et al. (2014). A 3D map of the human genome at kilobase resolution reveals principles of chromatin looping. Cell 159, 1665–1680.

Rubio, E.D., Reiss, D.J., Welcsh, P.L., Disteche, C.M., Filippova, G.N., Baliga, N.S., Aebersold, R., Ranish, J.A., and Krumm, A. (2008). CTCF physically links cohesin to chromatin. Proc. Natl. Acad. Sci. 105, 8309–8314.

Ruiz-Velasco, M., and Zaugg, J.B. (2017). Structure meets function: How chromatin organisation conveys functionality. Curr. Opin. Syst. Biol. 129–136.

Sanborn, A.L., Rao, S.S.P., Huang, S.-C., Durand, N.C., Huntley, M.H., Jewett, A.I., Bochkov, I.D., Chinnappan, D., Cutkosky, A., Li, J., et al. (2015). Chromatin extrusion explains key features of loop and domain formation in wild-type and engineered genomes. Proc. Natl. Acad. Sci. 112, E6456–E6465.

Schmidt, D., Wilson, M.D., Spyrou, C., Brown, G.D., Hadfield, J., and Odom, D.T. (2009). ChIP-seq: Using high-throughput sequencing to discover protein–DNA interactions. Methods 48, 240–248.

Schmidt, D., Wilson, M.D., Ballester, B., Schwalie, P.C., Brown, G.D., Marshall, A., Kutter, C., Watt, S., Martinez-Jimenez, C.P., Mackay, S., et al. (2010). Five-Vertebrate ChIP-seq Reveals the Evolutionary Dynamics of Transcription Factor Binding. Science (80-.). 328, 1036–1040.

Schmidt, D., Schwalie, P.C., Wilson, M.D., Ballester, B., Gonçalves, A., Kutter, C., Brown, G.D., Marshall, A., Flicek, P., and Odom, D.T. (2012). Waves of retrotransposon expansion remodel genome organization and CTCF binding in multiple mammalian lineages. Cell 148, 335–348.

Schoenfelder, S., Furlan-Magaril, M., Mifsud, B., Tavares-Cadete, F., Sugar, R., Javierre, B.-M., Nagano, T., Katsman, Y., Sakthidevi, M., Wingett, S.W., et al. (2015). The pluripotent regulatory circuitry connecting promoters to their long-range interacting elements. Genome Res. 25, 582–597.

Schwalie, P.C., Ward, M.C., Cain, C.E., Faure, A.J., Gilad, Y., Odom, D.T., and Flicek, P. (2013). Co-binding by YY1 identifies the transcriptionally active, highly conserved set of CTCF-bound regions in primate genomes. Genome Biol. 14, R148.

Sofueva, S., Yaffe, E., Chan, W.-C., Georgopoulou, D., Vietri Rudan, M., Mira-Bontenbal, H., Pollard, S.M., Schroth, G.P., Tanay, A., and Hadjur, S. (2013). Cohesin-mediated interactions organize chromosomal domain architecture. EMBO J. 32, 3119–3129.

Stedman, W., Kang, H., Lin, S., Kissil, J.L., Bartolomei, M.S., and Lieberman, P.M. (2008). Cohesins localize with CTCF at the KSHV latency control region and at cellular c-myc and H19/Igf2 insulators. EMBO J. 27, 654–666.

Sundaram, V., Cheng, Y., Ma, Z., Li, D., Xing, X., Edge, P., Snyder, M.P., and Wang, T. (2014). Widespread contribution of transposable elements to the innovation of gene regulatory networks. Genome Res. 24, 1963–1976.

Symmons, O., Uslu, V.V., Tsujimura, T., Ruf, S., Nassari, S., Schwarzer, W., Ettwiller, L., and Spitz, F. (2014). Functional and topological characteristics of mammalian regulatory domains. Genome Res. 24, 390–400.

Thybert, D., Roller, M., Navarro, F.C.P., Fiddes, I., Streeter, I., Feig, C., Martin-Galvez, D., Kolmogorov, M., Janoušek, V., Akanni, W., et al. (2018). Repeat associated mechanisms of genome evolution and function revealed by the Mus caroli and Mus pahari genomes. Genome Res. 28, 448–459.

Vietri Rudan, M., Barrington, C., Henderson, S., Ernst, C., Odom, D.T., Tanay, A., and Hadjur, S. (2015). Comparative Hi-C Reveals that CTCF Underlies Evolution of Chromosomal Domain Architecture. Cell Rep. 10, 1297–1309.

Wendt, K.S., Yoshida, K., Itoh, T., Bando, M., Koch, B., Schirghuber, E., Tsutsumi, S., Nagae, G., Ishihara, K., Mishiro, T., et al. (2008). Cohesin mediates transcriptional insulation by CCCTC-binding factor. Nature 451, 796–801.

Wong, E.S., Thybert, D., Schmitt, B.M., Stefflova, K., Odom, D.T., and Flicek, P. (2015). Decoupling of evolutionary changes in transcription factor binding and gene expression in mammals. Genome Res. 25, 167–178.

Xiao, T., Wallace, J., and Felsenfeld, G. (2011). Specific Sites in the C Terminus of CTCF Interact with the SA2 Subunit of the Cohesin Complex and Are Required for Cohesin-Dependent Insulation Activity. Mol. Cell. Biol. 31, 2174–2183.

Zhang, Y., Liu, T., Meyer, C.A., Eeckhoute, J., Johnson, D.S., Bernstein, B.E., Nussbaum, C., Myers, R.M., Brown, M., Li, W., et al. (2008). Model-based Analysis of ChIP-Seq (MACS). Genome Biol. 9, R137.

Zuin, J., Dixon, J.R., van der Reijden, M.I.J.A., Ye, Z., Kolovos, P., Brouwer, R.W.W., van de Corput, M.P.C., van de Werken, H.J.G., Knoch, T.A., van IJcken, W.F.J., et al. (2014). Cohesin and CTCF differentially affect chromatin architecture and gene expression in human cells. Proc. Natl. Acad. Sci. 111, 996–1001.

